# Synaptotagmin-like protein 2a regulates lumen formation via Weibel-Palade body apical secretion of angiopoietin-2 during angiogenesis

**DOI:** 10.1101/2021.02.15.431296

**Authors:** Caitlin R. Francis, Shea Claflin, Erich J. Kushner

**Author notes:** Author for correspondence: Erich J. Kushner, University of Denver, Department of Biological Sciences Denver, CO 80210.

## Abstract

**Objective:** Vascular lumen formation requires the redistribution of intracellular proteins to instruct apico-basal polarity, thereby enforcing maturation of both luminal and basal domains. In the absence of proper apical signaling, lumen formation can be distorted leading to lumen collapse and cessation of blood flow. Synaptotagmin-like protein-2a (Slp2a) has been implicated in apical membrane signaling; however, the role of Slp2a in vascular lumen formation has never been assessed.

**Approach and Results:** Our results demonstrate that Slp2a is required for vascular lumen formation. Using a 3- dimensional sprouting assay, sub-cellular imaging, and zebrafish blood vessel development we establish that Slp2a resides at the apical membrane acting as a tether for Rab27a that decorates Weibel-Palade bodies (WPBs). Unique to endothelial tissue, we show that Slp2a regulates exocytic activity of WPBs, thus regulating release of WPB contents into the luminal space during angiogenesis. Angiopoietin-2 is a Tie-2 receptor ligand that is selectively released from WPB secretory granules. We identify a critical role for angiopoietin-2 in regulating endothelial lumenization and show that in the absence of Slp2a, WPB contents cannot fuse with the apical membrane. This disrupts the release of angiopoietin-2 and blocks Tie-2 signaling necessary for proper lumen formation.

**Conclusions:** Our results demonstrate a novel requirement of Slp2a for vascular lumen formation. Moreover, we show that Slp2a is required for the exocytic release of WPB secretory granule cargo during vascular lumen development, and thus is a core upstream component of the WPB secretory pathway. Furthermore, we provide evidence that WPB-housed angiopoietin-2 is required for vascular lumen formation.

**HIGHLIGHTS:**

- Synaptotagmin-like protein-2a (Slp2a) is required for vascular lumen formation via its interaction with Rab27a and Weibel Palade Body secretory granules.
- Slp2a is recruited to the apical membrane where it regulates secretion of Weibel Palade Body components into the luminal space.
- In the absence of Slp2a, Weibel Palade Body-housed angiopoietin-2 ligand cannot be exocytosed, this impedes activation of Tie-2 signaling required for lumen biogenesis.
- Knockout of Slp2a or Tie-2 in zebrafish blunts the formation of vascular lumens during angiogenic development.

## INTRODUCTION

During development, new blood vessels emerge from pre-existing vasculature, a process termed angiogenesis [1-3]. During this time, endothelial cells (ECs) form a hollow opening, or central lumen. Vascular lumen formation can be roughly broken into three phases: 1) formation of a common cell-cell interface; 2) establishment of an apical membrane initiation site (AMIS) at the specific cell-cell interface promoting membrane deadhesion; and 3) lumen expansion. First, cadherin and integrin binding provide the initial cues for apical-basal polarity signaling in ECs [4, 5]. Thereafter, cell-cell adhesions localize laterally to allow for separation between neighboring cells [6, 7]. Concurrently, the AMIS located on the luminal membrane serves as a hub for asymmetric intracellular protein delivery to the maturing apical membrane. These AMIS trafficking events are responsible for delivering factors that cause deadhesion of opposing cell membranes as well as substantial cell shape changes leading to lumen cavity enlargement during angiogenesis [8-10]. For example, trafficking of sialomucin-laden glycoproteins, such as podocalyxin and CD34, to the apical membrane are required for lumen formation across multiple developmental models [8, 9, 11]. Precise trafficking of proteins to the maturing apical membrane are paramount to its biogenesis; however, what factors are involved in regulating trafficking during this critical period of vascular lumen formation are incompletely understood.

Synaptotagmin-like protein 2a (Slp2a), also called exophilin-4, is a phospholipid binding protein with high affinity for the apically enriched phospholipid, phosphotidylinositol (4,5) bisphosphate (PIP_2_) [12]. Characteristic of synaptotagmin family members, Slp2a interacts with phospholipids via its tandem C2 domains, C2A and C2B. Additionally, Slp2a’s Rab-binding domain provides interactions with Rab GTPases towing specific vesicle populations [13]. Existing evidence, based largely on studies in epithelia and melanocytes, indicates Slp2a principally binds Rab27a [14-17]. In endothelial cells Rab27a has been reported to decorate Weibel-Palade Bodies (WPBs), a prothrombotic secretory granule, and negatively regulate its exocytic activity [18]. Rab27a has also been shown to influence recycling of vascular endothelial growth factor receptor 1 (VEGFR1) [19]. In epithelium, Slp2a has been reported to tether podocalyxin-rich vesicles via Rab27a binding in cooperation with its family member synaptotagmin-like protein 4a (Slp4a) to promote lumen formation [15]. Slp2a has yet to be investigated in any aspect of blood vessel development.

In this report, our aim was to characterize Slp2a’s function during sprouting angiogenesis, with emphasis on its putative role in lumen formation. Our results demonstrate that Slp2a is required for vascular lumen formation in developing endothelial sprouts. We determine that in ECs, Slp2a is resident at the apical membrane during lumen initiation and expansion downstream of PIP_2_ lipid binding. Interestingly, deletion of the PIP_2_ interacting domains localized Slp2a exclusively to WPBs. We show that Slp2a is one of the most upstream components required for WPB secretion. Mechanistically, we determine that loss of Slp2a impedes angiopoietin-2 (Ang-2) secretion resulting in inhibited Tie-2 autocrine signaling, preventing lumen formation. Overall, our results demonstrate a novel role of Slp2a in regulating exocytic trafficking at the apical membrane and, in doing so, controlling Ang-2 release during blood vessel lumen formation.

## MATERIALS AND METHODS

Additional experimental procedures and a list of used materials is included in the Data Supplement. The authors will make their raw data, analytic methods, and study materials available to other researchers upon written request.

### Cell Culture

Pooled Human umbilical vein endothelial cells (HUVECs) were purchased from PromoCell and cultured in proprietary media (PromoCell) for 2-5 passages. For experiments glass-bottomed imaging dishes were exposed to deep UV light for 6 minutes and coated with Poly-D-Lysine (ThermoFisher) for a minimum of 20 minutes. Small interfering RNA (ThermoFisher) was introduced into primary HUVEC using the Neon® transfection system (ThermoFisher). Scramble, Slp2a, Rab27a, and Slp4a siRNAs were purchased from (ThermoFisher) and resuspended to a 10µM stock concentration and used at 0.5 µM. Normal human lung fibroblasts (NHLFs, Lonza) and HEK-A (ThermoFisher) were maintained in Dulbeccos Modified Medium (DMEM) supplemented with 10% fetal bovine serum and pen/strep antibiotics. Both NHLFs and HEKs were used up to 15 passages. All cells were maintained in a humidified incubator at 37°C and 5% CO_2_.

Phorbol myristate acetate (PMA) or histamine (Sigma) was used to induce secretion of WPB components. To achieve this, cells were serum-starved for 6 hours and treated with a final concentration of 100ng/mL PMA or 100 µM histamine for 15 minutes. Cells were then washed with phosphate buffered saline (PBS) and fixed promptly in 4% paraformaldehyde (PFA). For Tie- 2 inhibition, cells were treated with BAY-826 (TOCRIS) at a final concentration of 1.3 nM for 1-3 days during sprouting.

### Sprouting Angiogenesis Assay

Fibrin-bead assay was performed as reported by Nakatsu et al. 2007 [20]. Briefly, HUVECs were coated onto microcarrier beads (Amersham) and plated overnight. SiRNA-treatment or viral transduction was performed the same day the beads were coated. The following day, the EC-covered microbeads were embedded in a fibrin matrix. Once the clot was formed media was overlaid along with 100,000 NHLFs. Media was changed daily along with monitoring of sprout development.

### Plasmid Constructs

The following constructs were procured for the study: pGEX4T-1 (gift from Fernando Martin-Belmonte; Addgene plasmid #40059); pmCherry-C1 hSlp2-a (gift from Fernando Martin-Belmonte; Addgene plasmid #40056); pEGFP-C1 hSlp2-a C2AB (gift from Fernando Martin-Belmonte; Addgene plasmid #40051); pEGFP-C1 hSlp2-a del.SHD (gift from Fernando Martin-Belmonte; Addgene plasmid #40050); pEGFP-C1 hSlp4-a (gift from Fernando Martin-Belmonte; Addgene plasmid #40032); GFP-Rab27A (gift from William Gahl; Addgene plasmid #89237); pro-vWF-GFP/mCherry (gift from Tom Carter), and pCMV6-Angpt2 (Origene MR207970).

### Lentivirus and Adenovirus Generation and Transduction

Please see data supplement.

### Immunofluorescence and Microscopy

Prior to seeding cells, coverslips were treated with poly-D Lysine for approximately 20 minutes and washed 2 times with PBS. HUVECs were fixed with 4% PFA for 7 minutes. ECs were then washed three times with PBS and permeabilized with 0.5% Triton-X (Sigma) for 10 minutes. After permeabilization, cells were washed three times with PBS. ECs were then blocked with 2% bovine serum albumin (BSA) for 30 minutes. Once blocked, primary antibodies were incubated for approximately 4-24 hours. Thereafter, primary antibodies were removed and the cells were washed 3 times with PBS. Secondary antibody with 2% BSA were added and incubated for approximately 1-2 hours, washed 3 times with PBS and mounted on a slide for imaging.

For imaging the fibrin-bead assay, first fibroblasts were removed from the clot with a 1- minute trypsin incubation. Following incubation, the trypsin was neutralized with DMEM contain 10% BSA, washed 3 times with PBS, and fixed using 4% paraformaldehyde for 40 minutes. After fixation, the clot was washed 3 times with PBS, permeabilized with 0.5% Triton-X for 2 hours and then blocked with 2% BSA for 1 hour prior to overnight incubation with primary antibodies. The following day, primary antibodies were removed and the clot was washed 5 times with PBS and secondary antibody was added with 2% BSA and incubated overnight. Prior to imaging the clot was washed 5 times with PBS. All primary and secondary antibodies are listed in the Data Supplement. Images were taken on a Nikon Eclipse Ti inverted microscope equipped with a CSU-X1 Yokogawa spinning disk field scanning confocal system and a Hamamatusu EM-CCD digital camera. Cell culture images were captured using a Nikon Plan Apo 60x NA 1.40 oil objective using Olympus type F immersion oil NA 1.518. All images were processed using ImageJ (FIJI).

### Immunoblotting & Protein Pull-Down

Please see data supplement.

### Zebrafish Experiments

Please see data supplement.

### Statistical Analysis

Experiments were repeated a minimum of three times. Statistical analysis and graphing was performed using GraphPad Prism. Statistical significance was assessed with a student’s unpaired t-test for a two-group comparison. Multiple group comparisons were carried out using a one-way analysis of variance (ANOVA) followed by a Dunnett multiple comparisons test. Data was scrutinized for normality using Kolmogorov-Smirnov (K-S) test. Zebrafish gender distribution was not adjusted as sex determination did not occur at the stage of development in which the specimens were assayed. Statistical significance set a priori at p<0.05.

## RESULTS

### Slp2a is apically localized and required for lumen formation *in vitro*

We first sought to determine Slp2a’s localization and function *in vitro* given its spatial organization in vascular sprouting was not previously characterized. To do so, we transduced an mCherry-tagged Slp2a virus into primary ECs in a 3-dimensional (3D)-sprouting assay that closely mimics *in vivo* sprouting angiogenesis [20, 21] (**Fig. 1A**). Here, ECs are coated onto micro-carrier beads, embedded in a fibrin matrix and allowed to sprout for 4-5 days. Transduced ECs were stained for the junctional marker vascular-endothelial (VE)-cadherin to identify cell-cell interfaces as well as moesin and podocalyxin to delineate the apical membrane (**Fig. 1B**). Prior to lumen formation, Slp2a heavily colocalized with all three proteins at cell-cell junctions (**Fig. S1A**). However, in sprouts with an established lumen opening, Slp2a was only located on the apical membrane, strongly colocalizing with apical markers moesin and podocalyxin, but distinct from VE-cadherin at cell-cell interfaces (**Fig. 1B**). These results indicate that Slp2a is preferentially localized to the apical membrane prior to and throughout lumen formation, consistent with reports in epithelial tissues [15]. Slp2a’s C2AB domains are purported to bind PIP_2_, a well-known apical lipid species [13, 15, 22]. We confirmed this interaction in ECs using a PIP_2_ biosensor (PH-GFP). [23] in which Slp2a and PH-GFP dynamically colocalized at junctions in 2D culture (**Fig. S1B,C**). Overall, these results demonstrate that Slp2a is resident at the apical membrane downstream of binding to PIP_2_.

**Figure 1.**
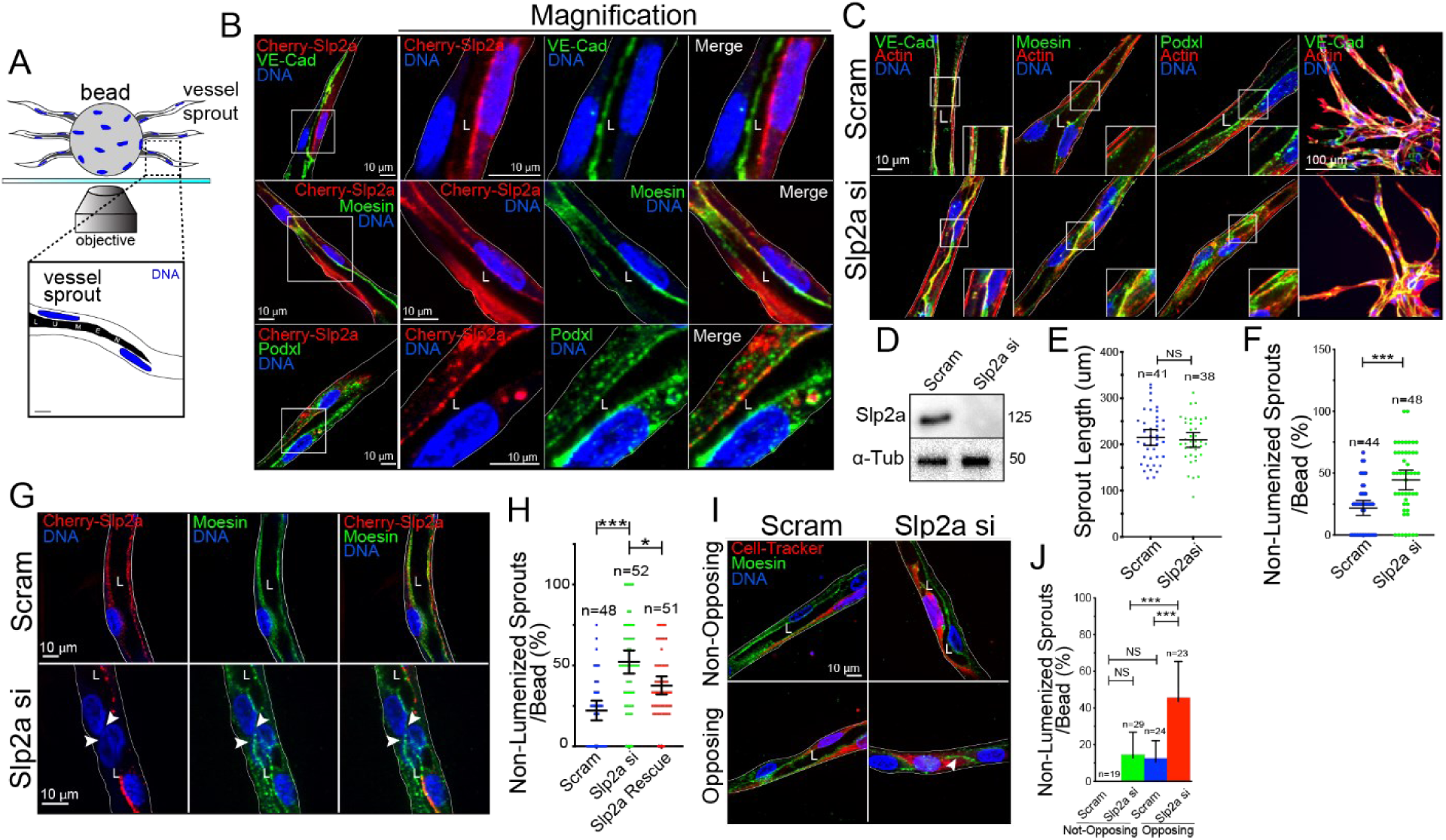
Slp2a is an apically localized protein required for sprout formation. **A,** Cartoon model of 3-dimensional sprouting assay denoting imaging setup. **B,** Localization of transduced mCherry (Cherry)-Slp2a during lumen formation in sprouts stained for VE-Cadherin (VE-Cad), moesin, and podocalyxin (Podxl). **C**, Images of Slp2a siRNA (si) knockdown and scramble (scram) control sprouts stained for indicated proteins. **D,** Confirmation of Slp2a siRNA-mediated knockdown by western blot probed for Slp2a and alpha-tubulin (α-Tub). n=3. **E,** Quantification of sprout length for indicated groups. **F,** Quantification of non-lumenized sprouts between indicated groups. **G,** Mosaic rescue experiment in which cells were treated with indicated siRNAs and transduced with Cherry- Slp2a (red). Arrows indicate a lack of lumen in addition to a lack of Cherry-Slp2a expression. **H,** Quantification of percent non-lumenized sprouts between indicated groups. **I,** Mosaic knockdown experiment in which cells were treated with Slp2a siRNA (red) and then mixed with scramble-treated cells (non-fluorescent) and then challenged to sprout. Top row depicts non-opposing siRNA-treated cells. Bottom row depicts opposing siRNA-treated cells. Arrow denotes a lack of lumen. **J,** Quantification of mosaic KD sprouts with percent non-lumenized sprouts. All experiments used human umbilical vein endothelial cells. In all panels L denotes lumen; white box denotes magnification; white lines denotes exterior of sprout; values are means +/- SEM; n= individual sprouts across three experimental replicates; significance: *P<0.05, ***P<0.0005, NS=Not Significant. Statistical significance was assessed with an unpaired t-test or a 1-way ANOVA followed by a Dunnett multiple comparisons test.

We next asked if Slp2a played a role in angiogenic sprouting and lumen formation via loss of function using small-interfering (si)-RNA knockdown. Morphologically, Slp2a knockdown did not alter migration programs as sprout lengths were unaffected (**Fig. 1E**); however, the sprouts were visibly thinner in appearance compared with controls (**Fig. 1C,D; S1D**). To investigate this phenotype, we quantified the percentage of non-lumenized sprouts (sprouts with no discernable, or contiguous, lumen cavity). We determined that loss of Slp2a significantly increased the percentage of non-lumenized sprouts compared with controls (**Fig. 1F**). We next performed a rescue experiment by mosaically overexpressing mCherry-Slp2a on a Slp2a knockdown background to further examine if Slp2a deficiency was underlying the lack of lumen formation. ECs expressing mCherry-Slp2a exhibited a significant increase in lumen formation compared with non-transduced controls (**Fig. 1G,H**). In a similar approach, we tested if loss of Slp2a was cell autonomous in terms of its impact on lumenogenesis. To do so, we knocked down Slp2a in a population of ECs that were then labeled with red CellTracker. This population was mixed with scramble siRNA-treated ECs and challenged to sprout. We observed that in Slp2a knockdown ECs opposite wild-type (WT) a lumen opening was maintained, albeit small; while two opposing knockdown ECs failed to create a luminal cavity (**Fig. 1I,J**). These results suggest that Slp2a is cell autonomous and required for lumen formation *in vitro*.

### Slp2a interacts with Weibel-Palade Bodies

To better understand the mechanism(s) by which a Slp2a deficiency results in lumen defects, we employed two Slp2a domain mutants: 1) a deletion of the PIP_2_ binding C2AB domains (Slp2a-ΔC2AB); and 2) expression of only the C2AB domains (Slp2a-C2AB) (**Fig. 2A**). In a mosaic rescue assay, both mCherry-Slp2a-ΔC2AB and mCherry-Slp2a-C2AB mutants were transduced into sprouts on a Slp2a siRNA knockdown background. Neither mutant proved capable of rescuing lumen abnormalities (**Fig. 2B-D**), suggesting both domains are required for Slp2a to function properly during vascular lumen formation. Upon further inspection, the Slp2a- C2AB mutant localized largely to the apical membrane similar to WT Slp2a in lumenized sprouts. Conversely, the mCherry-Slp2a-ΔC2AB mutant no longer localized to the apical membrane, but decorated rod-like puncta that strongly colocalized with von Willebrand Factor (vWF), a well-established WPB marker (**Fig. 2E; S2A**) [18]. We next determined if the Slp2a-ΔC2AB -decorated WPBs showed any localization preference during the lumenization process. The Slp2a-ΔC2AB- decorated WPBs exhibited a heightened cytoplasmic distribution in non-lumenized sprouts. However, in sprouts with a defined lumen, the Slp2a-ΔC2AB-decorated WPBs, preferentially localized to the apical membrane (**Fig. 2F,G; S2B,C**). Taken together, in the absence of membrane binding, Slp2a’s default localization is on WPBs that are being actively transported to the apical membrane during lumenogenesis.

**Figure 2.**
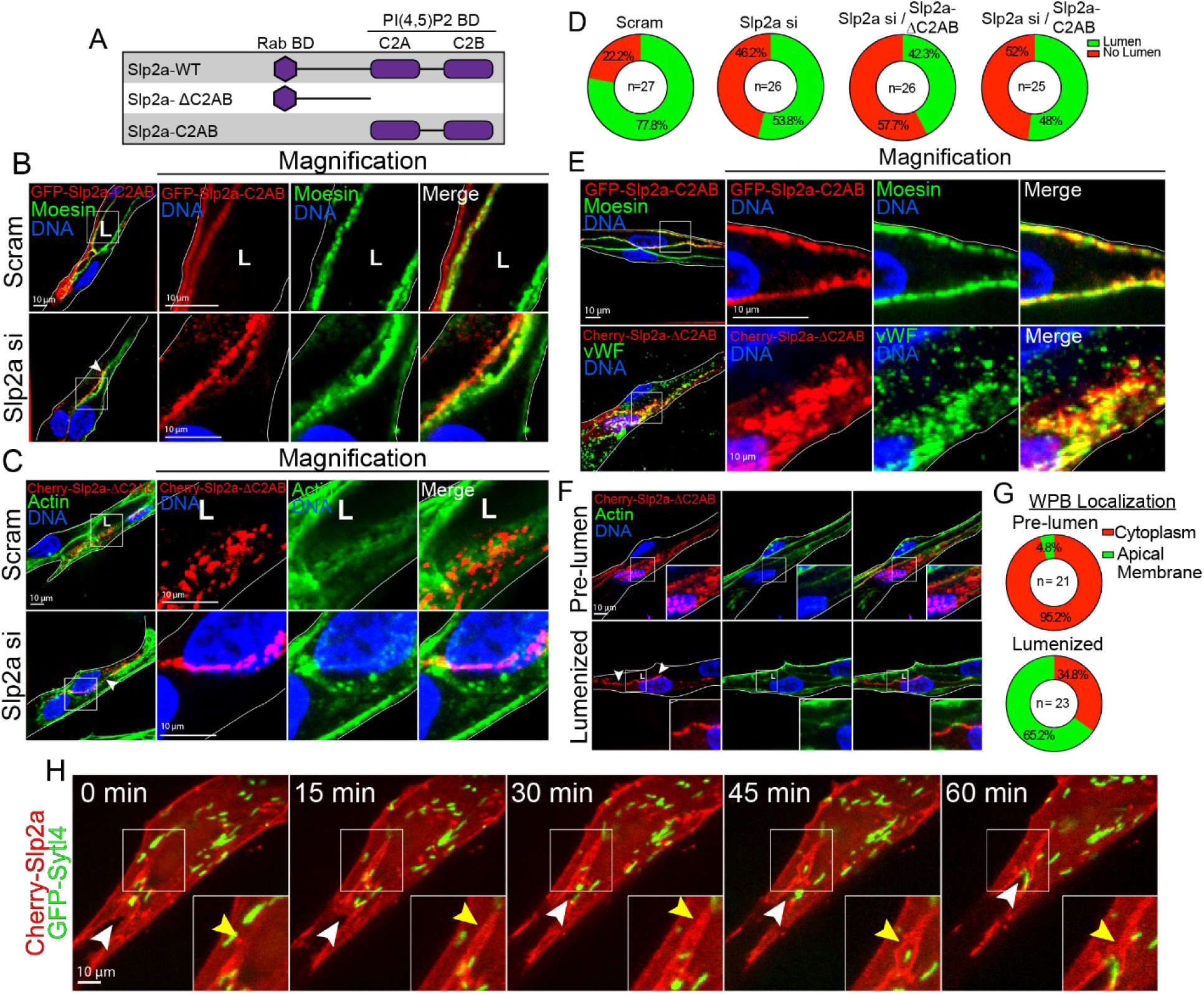
Slp2a lacking C2 domains localizes to Weibel-Palade bodies. **A,** Cartoon model of Slp2a domains and mutants used for experimentation. Slp2a-ΔC2AB lacks two phospholipid binding C2 domains. Slp2a-C2AB mutant lacks the Rab-binding domain as well as residues linking it to the C2AB domains. **B,** GFP-Slp2a-C2AB expressing in both scramble (scram) and Slp2a siRNA(si) knockdown groups and stained for indicated proteins. Arrow indicates lack of lumen. **C,** MCherry(Cherry)-Slp2a-ΔC2AB expressing in both scramble and Slp2a siRNA knockdown groups and stained for indicated proteins. Arrow indicates lack of lumen. **D,** Quantification of lumen formation of individual sprouts. Cells were treated with scramble or Slp2a siRNA and then infected with indicated constructs. Green represents lumen formation and red represents non-lumenized sprouts. N-value represents individual sprouts across three experimental replicates. **E,** GFP-Slp2a-C2AB and Cherry-Slp2a-ΔC2AB expressing in sprouts stained for moesin and with Weibel-Palade body (WPB) marker, von Willebrand factor (vWF). **F,** Localization of Cherry-Slp2a-ΔC2AB prior to lumen opening (pre-lumen, top panels) and after lumen opening (lumenized, bottom panels). Arrows indicate heavy localization at the apical membrane. **G,** Quantification of Cherry-Slp2a-ΔC2AB localization pre-lumen and during lumenogenesis (lumenized). N-value represents individual sprouts across three experimental replicates. **H,** Live imaging of mCherry-Slp2a-WT and GFP-Slp4a-WT. Yellow arrow identifies future lumen expansion and white arrow indicates open lumen. All experiments use human umbilical vein endothelial cells. In all panels L denotes lumen; white box denotes magnification.

Slp4a has previously been reported to interact with Slp2a in epithelial cells [15]. In endothelium, Slp4a is reported to decorate WPBs, but has not yet been functionally linked to Slp2a [24]. To determine if Slp4a interacted with Slp2a we expressed both family members at the same time. Slp2a and Slp4a exhibited disparate localization patterns; Slp2a maintained its localization at the apical membrane, while Slp4a resided on WPBs (**Fig. S3A,B**). Similarly, live-imaging of Slp2a and Slp4a in 3D sprouts revealed that areas of lumen formation were decorated by Slp2a, while Slp4a-positive WPBs trafficked to the apical membrane, presumably for exocytosis of WPB secretory granules into the luminal space (**Fig. 2H**). Given Slp2a and Slp4a’s previous association, we wanted to determine if Slp4a played a role in lumen formation. Knockdown of Slp4a did not affect sprouting or lumen formation parameters (**Fig. S3C,D**). These data suggest that Slp2a is distinct from Slp4a in its localization and role in lumen formation.

### Slp2a binds Rab27a resident on WPBs

Rab27a has been shown to directly bind Slp2a in other systems as well as in ECs [13, 15, 18]. To test if this was true in our model, we overexpressed a GFP-tagged Rab27a construct. Overexpression of WT Rab27a in 2D culture produced colocalization with WT Slp2a at discrete puncta, while Slp2a was also located at the membrane (**Fig. 3A**). Slp2a-ΔC2AB mutant over- expression in 2D strongly colocalized with Rab27a puncta only (**Fig. 3A**). In 3D sprouts, Rab27a and Slp2a did not show similar localization patterns. Slp2a localized solely to the apical membrane while Rab27a was on WPB puncta and, to some extent, on the apical membrane (**Fig. 3B**). However, expression of Slp2a-ΔC2AB mutant exhibited strong colocalization with Rab27a puncta that were localized to WPBs in sprouts (**Fig. 3B; S4A**). We confirmed this direct interaction via immunoprecipitation using a GST-tagged Slp2a as bait and detected Rab27a binding (**Fig. 3C**). Next, we performed a mitochondrial mis-targeting assay. Here, a mitochondrial-targeting sequence (Tom20) [25] was added to the N-terminal GFP-tag to unnaturally anchor Rab27a to the outer mitochondrial membrane (**Fig. 3D**). This allowed us to visualize what proteins or complexes were ‘pulled along’ with Rab27a to the mitochondria as an intracellular readout for binding interactions. Wild-type Slp2a moderately and the Slp2a-ΔC2AB mutant strongly localized to the mitochondria in ECs expressing Tom20-GFP-Rab27a, suggesting that Slp2a is binding Rab27a (**Fig. 3E**). To determine if this binding was dependent on Rab27a’s activation state, either an inactive form (GDP) or an active form (GTP), we performed the same experiment using a constitutively active (CA, Q78L) and a dominant negative (DN, L130P) Rab27a mutant [26]. Co-expression of Slp2a-ΔC2AB with Rab27a CA exhibited robust colocalization at the mitochondria, while expression of the Rab27a DN mutant abolished mitochondrial localization of Slp2a-ΔC2AB (**Fig. 3E**). Overall scoring of Slp2a localization between all above conditions clearly indicated that Slp2a binds Rab27a in a GTP-dependent fashion (**Fig. 3F**). Given Rab27a was located on WPBs, we also probed for vWF to determine if mis-localizing Rab27a also distorted the spatial distribution of WPB cargo. Mitochondrial-targeted Rab27a demonstrated a mixed phenotype: in some instances, WPBs were mislocalized to the mitochondria; however, in others, WPBs were not mis- targeted (**Fig. S4B,C**). Overall, this data suggests the Rab27a and Slp2a are robust binding partners.

**Figure 3.**
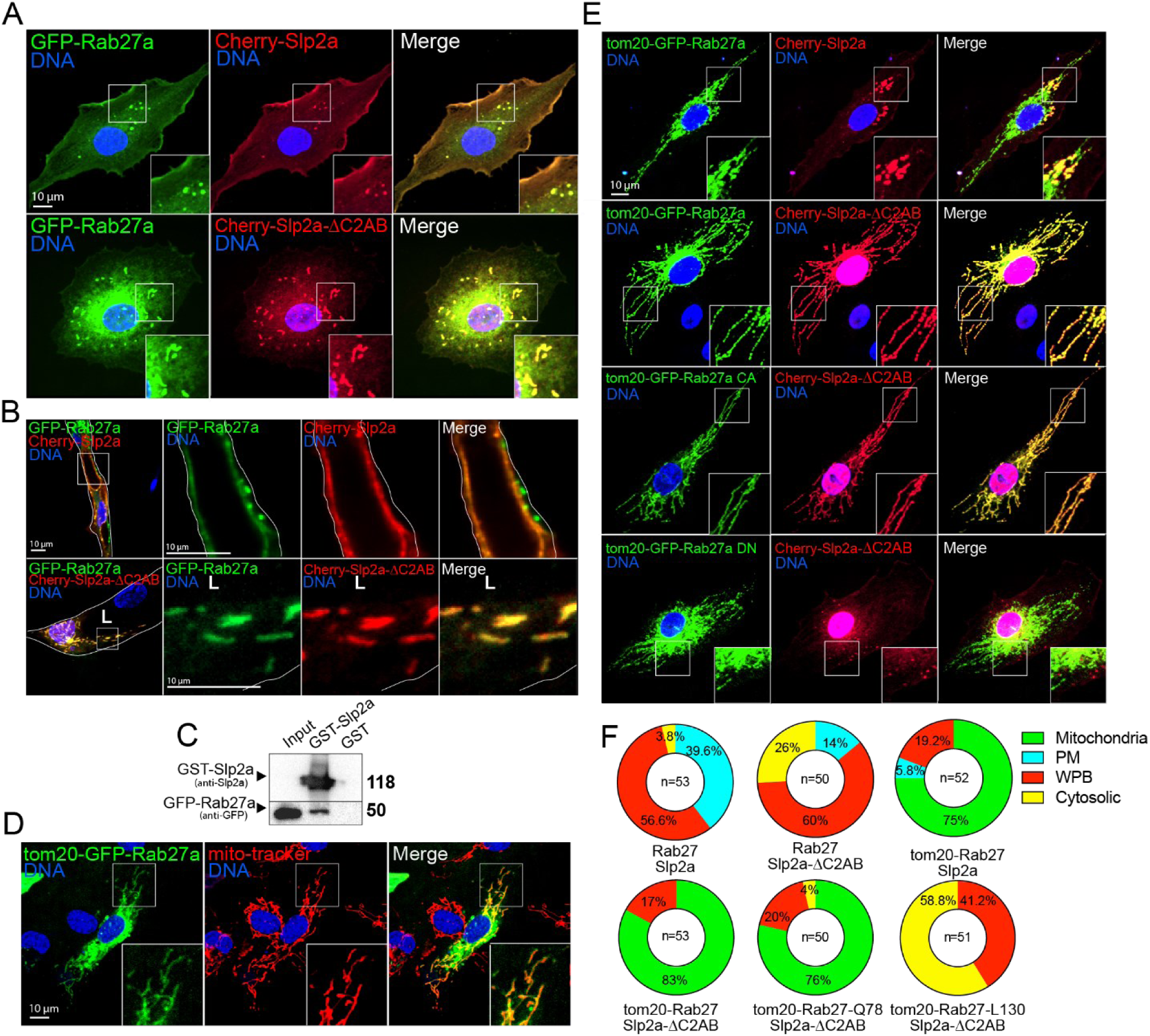
Slp2a binds Rab27a. **A,** 2-dimensional localization of mCherry(Cherry)-Slp2a and GFP-Rab27a in top panel. Bottom panel, localization of Cherry-Slp2a-ΔC2AB and GFP-Rab27a. **B,** Localization of Cherry-Slp2a and GFP-Rab27a in sprouts (top panels) and Cherry-Slp2a-ΔC2AB and GFP-Rab27a (bottom panels). **C,** Representative image of immunoprecipitation of GST-tagged Slp2a and GST (control) proteins used to probe for Rab27a binding. Image is one of three experimental replicates. **D,** Tom20-tagged GFP-Rab27a expressing cells also stained for mitochondria (Mito-tracker®). **E,** Tom20-GFP-Rab27a mis-localization experiments in 2D to test for binding interactions. Rab27a constitutively active (CA, Q78L) and dominant negative (DN, L130P) mutants were co-expressed with Cherry-Slp2a-ΔC2AB and Cherry-Slp2a. **F,** Quantification of localization of Cherry-Slp2a-ΔC2AB and mCherry-Slp2a in each of the 2D experiments presented in panel E. n=number of individual cells over three experimental repeats. All experiments use human umbilical vein endothelial cells. In all panels L denotes lumen; white box denotes magnification; white lines denote exterior of sprout.

Previous studies in non-endothelial tissues have reported that Rab27a transports podocalyxin, a negatively charged glycoprotein shown to be required for lumen formation [8, 9, 11, 17, 27]. Thus, Slp2a may be mediating podocalyxin transport by way of Rab27a. This association could potentially explain why loss of Slp2a results in lumen formation defects. To explore this, we determined the localization of both Rab27a and podocalyxin in lumenizing sprouts. Neither, Rab27a or vWF colocalized with podocalyxin, suggesting Rab27a is not interfacing with this protein during vascular lumenogenesis (**Fig. S4D**). Additionally, Rab27a knockdown did not affect podocalyxin localization; also suggesting that Rab27a does not transport podocalyxin in ECs (**Fig. S4E**).

### Slp2a regulates WPB exocytosis

Since Slp2a and Rab27a demonstrated direct binding, we next tested whether Rab27a was involved in lumen formation during angiogenic sprouting. In the 3D sprouting assay, siRNA knockdown of Rab27a did not affect lumen formation compared with a Slp2a knockdown (**Fig. 4A,B**). Interestingly, lumen diameter was significantly larger in the absence of Rab27a, whereas ablation of Slp2a in any condition abolished lumen formation resulting in significantly thinner sprouts (**Fig. 4C**). This data suggests that although Slp2a and Rab27a are bona fide binding partners, Rab27a does not negatively affect lumen formation during angiogenic sprouting.

**Figure 4.**
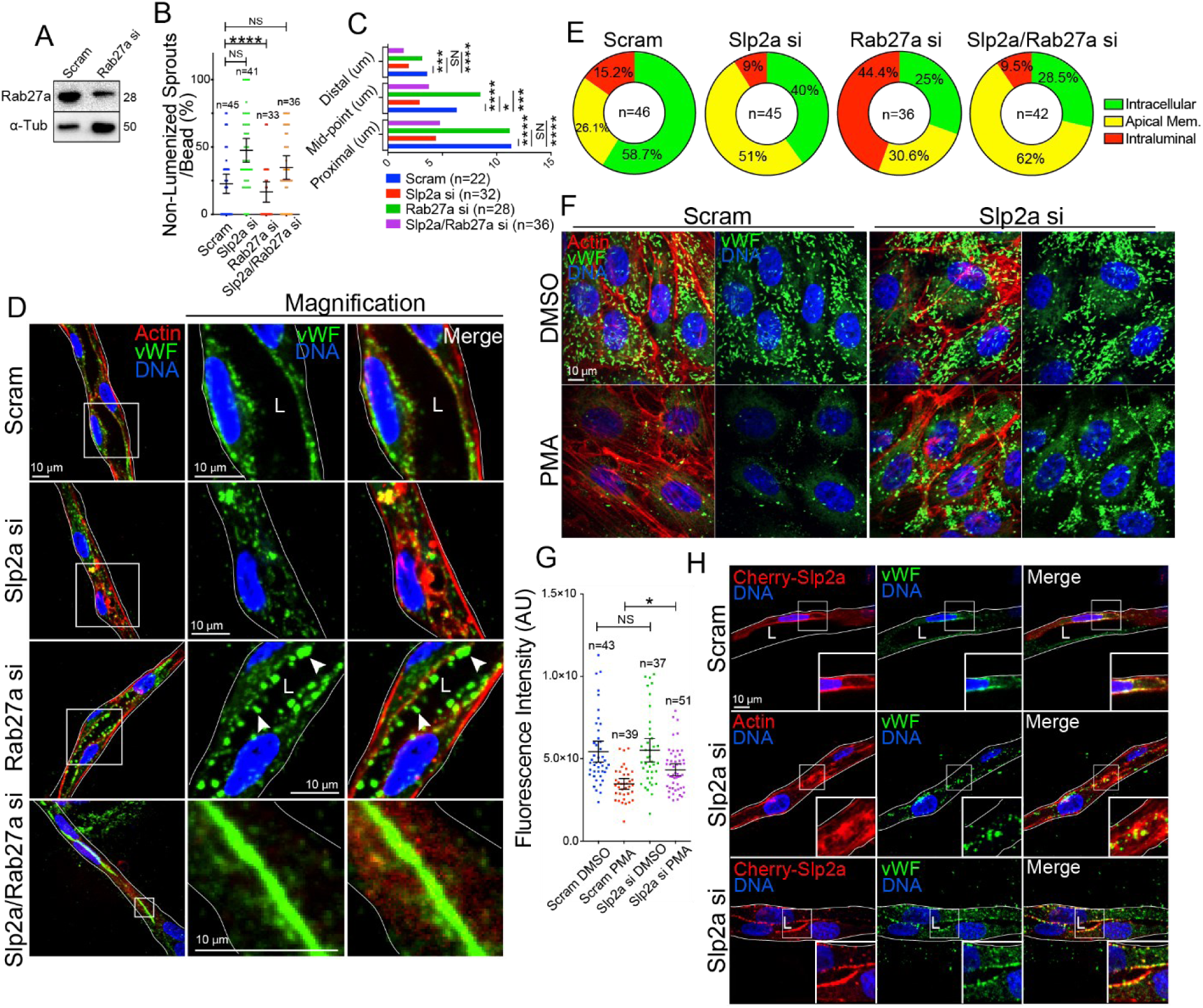
Slp2a is required for WPB exocytosis. **A,** Representative western blot confirmation of Rab27a siRNA (si) knockdown. n=3. **B,** Quantification of non-lumenized sprouts in indicated groups. n= individual sprouts over three experimental repeats. **C,** Quantification of lumen diameter at multiple locations within sprouts. Distances were measured proximally, at the mid- point, and distally from the bead. n=individual sprouts over three experimental repeats. **D,** Localization of Weibel-Palade body cargo von-Willebrand Factor (vWF), during lumen formation between siRNA-treated groups. Arrows indicate accumulation of vWF within the lumen. **E**, Quantification of vWF localization in indicated siRNA-treated groups. N= individual cells across three experimental repeats. **F,** Images of phorbol 12-myristate 13-acetate (PMA)- and vehicle (DMSO)-treated cells between indicated groups. **G,** Quantification of vWF fluorescent intensity between indicated conditions. n= individual cells across three experimental repeats. **H,** Mosaic rescue effect on vWF localization in sprouts between indicated groups. Cells were transduced with mCherry (Cherry)-Slp2a and treated with indicated siRNA. All experiments use human umbilical vein endothelial cells. AU= arbitrary unit. In all panels L denotes lumen; white box denotes magnification; white lines denote exterior of sprout; values are means +/- SEM; significance: *P<0.05, ***P<0.001, ****P<0.0001, NS=Not Significant. Statistical significance was assessed with a 1-way ANOVA followed by a Dunnett multiple comparisons test.

Given both Slp2a and Rab27a interact with WPBs, we next tested their respective roles in WPB-mediated exocytosis of vWF in 3D sprouts. As previously shown (**Fig. 1C-F**), loss of Slp2a resulted in a lack of lumen formation, thus there was little-to-no apical space for vWF to be secreted. As such, in Slp2a knockdown sprouts we observed vWF contained within the cytoplasm adjacent to sites of vacuolation (**Fig. 4D**). In this condition, vWF puncta did not accumulate at interior junctions, the presumptive sites of lumen expansion. By contrast, knockdown of Rab27a resulted in a robust secretion of vWF into the luminal cavity (**Fig. 4D**). This finding is in line with previous literature designating Rab27a as a negative regulator of WPB exocytosis [24], although, this has not been demonstrated in 3D sprouts. Double knockdown of Slp2a and Rab27a resulted in a dramatic accumulation of vWF at cell-cell junctions (**Fig. 4D,E**). As the loss of Slp2a abolished lumen formation, we could not ascertain if the vWF was able to be secreted into the lumen or was trapped in the subapical space. To address this, we performed the same experiment in 2D culture to track vWF secretion. First, we compared phorbol myristate acetate (PMA)-induced vWF secretion with and without Slp2a knockdown. Loss of Slp2a significantly reduced the ability of vWF to be secreted into the media compared to controls (**Fig. 4F,G**). Histamine-mediated release of vWF was also blunted in the absence of Slp2a compared with controls (Fig. S5A-C). To further test Slp2a’s involvement in vWF secretion, we performed a rescue experiment by overexpressing mCherry-Slp2a on a knockdown background. We observed that ECs overexpressing mCherry-Slp2a were capable of trafficking vWF to the apical membrane to a greater extent than non-transduced controls (**Fig. 4H**). Overall, these results indicate that Slp2a is likely an upstream regulator of WPB exocytosis.

We next aimed to understand if Slp2a or Rab27a affected each other’s localization in 3D sprouts. In other words, is there a dependency between Slp2a and Rab27a for localization to the apical membrane or on WPBs? Loss of Slp2a did not affect Rab27a or Slp4a’s localization to WPBs (Fig. S6A,B). Similarly, knockdown of Rab27a did not affect Slp2a localization to the apical membrane during lumen formation or Slp4a’s localization to WPBs (Fig. S6C,D). To also explore if knocking down either Slp2a or Slp4a altered each other’s expression levels, we probed for protein levels. Knockdown of Slp2a did not affect expression of Slp4a and visa versa (Fig. S6E). Similarly, overexpression of Slp2a-ΔC2AB did not affect levels of Slp4a on WPBs (Fig. S6F,G). These data indicate that Slp2a does not affect Rab27a and Slp4a’s ability to localize to WPBs. In addition, Slp2a localization to the apical membrane is not dependent on Rab27a.

### Secretion of Ang-2 is required for lumen formation

Given loss of Slp2a results in elevated non-lumenized sprouts and ablated WPB exocytosis, we postulated that vascular lumenization required the secretion of a WPB-housed factor(s) whose secretion was being controlled by Slp2a. Of the many proteins reported to be contained within WPBs, Ang-2 has been shown to have a proangiogenic effect in certain circumstances by differentially regulating Tie-2 signaling [28, 29]. To determine if Ang-2 was resident in the same WPB population that Slp2a decorated, we constructed an RFP and GFP-tagged version of Ang-2. Expression of Ang-2 demonstrated strong colocalization with vWF, Rab27a, and Slp2a-ΔC2AB positive WPBs in 2D culture (**Fig. 5A**). Here, we did observe some Slp2a-ΔC2AB puncta that did not colocalize with Ang-2, Ang-2 potentially being in non-WPB cytoplasmic granules (**Fig. 5B**). Interestingly, we also observed that in ECs containing WPBs, Ang-2 was packaged into dense WPB puncta; however, in ECs lacking WPBs Ang-2 was largely scattered throughout the cytoplasm in granular puncta (**Fig. 5B**). This phenotype was not observed in 3D sprouts (**Fig. 5C,D**), suggesting the 3D environment promotes Ang-2 trafficking via the WPB pathway to a greater extent than 2D culture. Next, we investigated when Ang-2 was being released during lumen formation. In sprouts actively forming a luminal surface there was elevated levels of Ang-2 localized to the apical membrane as compared to sprouts that already established a stable lumen cavity (Fig. 5E,F). In total, these data suggest that Ang-2 is housed in Slp2a-decorated WPBs and is targeted to the apical membrane during lumen formation.

**Figure 5.**
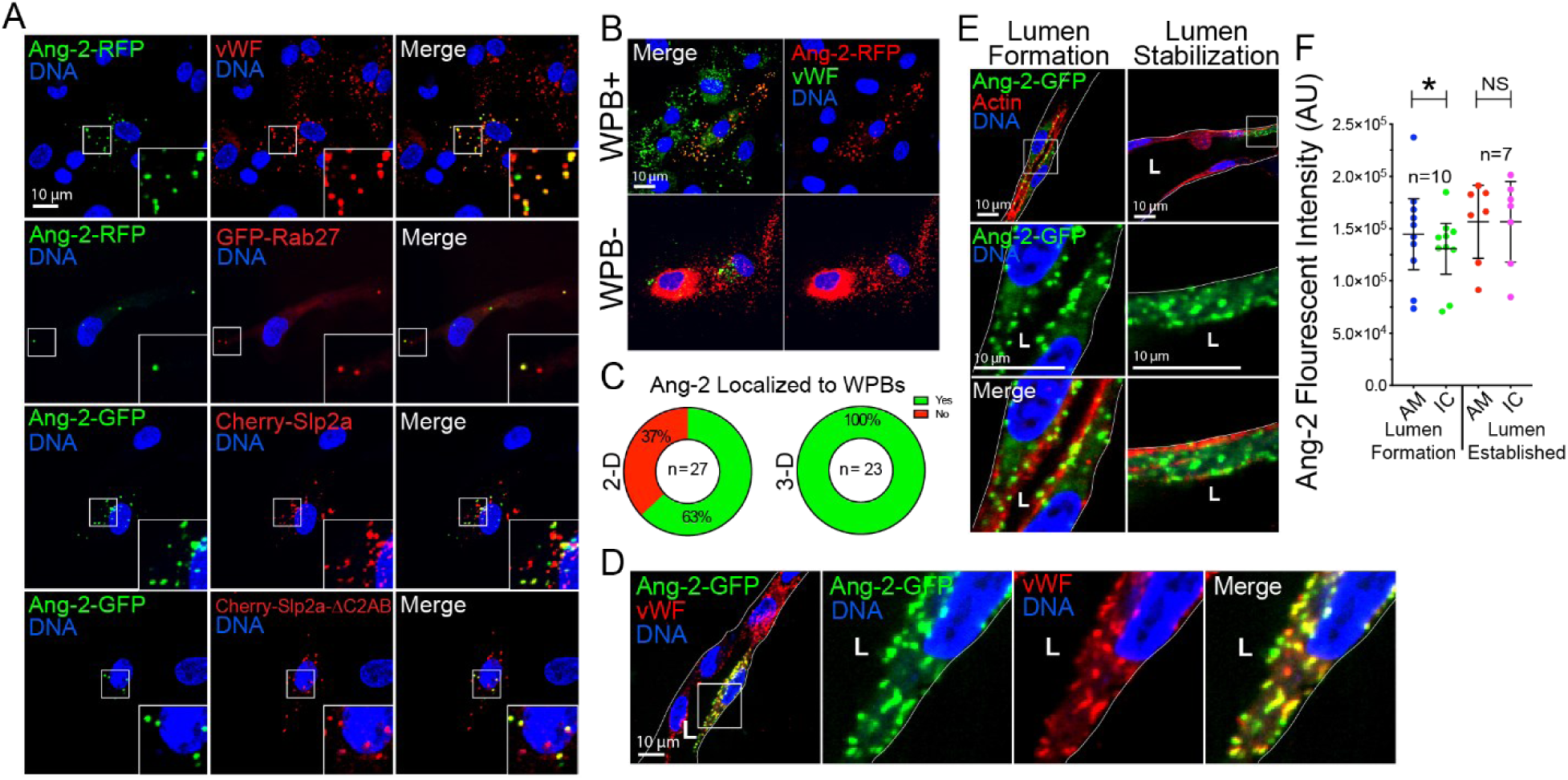
Angiopoietin-2 is housed within Slp2a-ΔC2AB positive WPBs. **A,** Angiopoietin-2 (Ang-2) colocalization experiments in 2-dimensional (2D) culture. Images show localization of Ang-2-RFP, GFP-Rab27a, von-Willebrand Factor (vWF), mCherry (Cherry)-Slp2a and Cherry- Slp2a-ΔC2AB. **B,** Ang-2 in cells with and without Weibel-Palade bodies (WPBs) denoted by vWF-positive staining. **C,** Quantification of Ang-2-GFP localization to WPBs between 2D culture and 3D sprouts. **D,** Ang-2-GFP and vWF localization in 3D sprouts. **E,** Ang-2-GFP localization at different time points during sprout development. The left panels are localization during the early stage of lumen formation and the right panels are after lumens are established. **F,** Quantification of Ang-2-GFP localization at different developmental time points. AU= arbitrary unit, AM= apical membrane and IC= intracellular. n=individual sprouts over three experimental repeats. All experiments use human umbilical vein endothelial cells. In all panels L denotes lumen; white box denotes magnification; white lines denote exterior of sprout; values are means +/- SEM; significance: *P<0.05, NS=Not Significant. Statistical significance was assessed with an unpaired t-test.

To determine if Ang-2 was required for lumen formation, we knocked down Ang-2 in 3D sprouts. Loss of Ang-2 phenocopied Slp2a knockdown in significantly elevating the percentage of non-lumenized sprouts (Fig. 6B,C**; S7A**). Depending on the context, Ang-2 has been shown to both activate Tie-2 signaling [30, 31] or act as an antagonist to Ang-1 limiting Tie-2 activation [29, 32]. Staining phosphorylated Tie-2 (pTie-2) revealed strong localization at the apical membrane and at cell-cell junctions in sprouts undergoing active lumen formation (**Fig. 6A**). Loss of Slp2a or Ang-2 significantly reduced pTie-2 activation at the apical membrane (**Fig. 6D**), indicating that Ang-2 is enforcing Tie-2 activation during lumen formation. To investigate if Tie-2 activation was necessary for lumen development we added the Tie-2 inhibitor Bay-826 on different days during lumen development (Fig. 6E,F**; S7B**). Tie-2 inhibition significantly increased the percentage of non-lumenized sprouts on day-1 and day-2 which coincide with the key stages of lumen formation (**Fig. 6G**). However, inhibiting Tie-2 activation on day-3 did not significantly impact lumen development, indicating Tie-2 activation is required for lumen formation, not maintenance. To ensure, Slp2a was controlling Ang-2 release, we assayed for secreted Ang-2 in the culture media. Knockdown of Slp2a reduced the amount of Ang-2 present in the media, suggesting blunted secretion (**Fig. 6H**). Likewise, we assayed the intracellular Ang-2 pool, reasoning that if Slp2a is diminishing secretion there would be increased intracellular retention. Indeed, loss of Slp2a resulted in higher intracellular Ang-2 compared with control (**Fig. 6I**). Overall, our results suggest that Slp2a regulates the release of Ang-2 which is necessary for lumen formation via activation of Tie-2 signaling.

**Figure 6.**
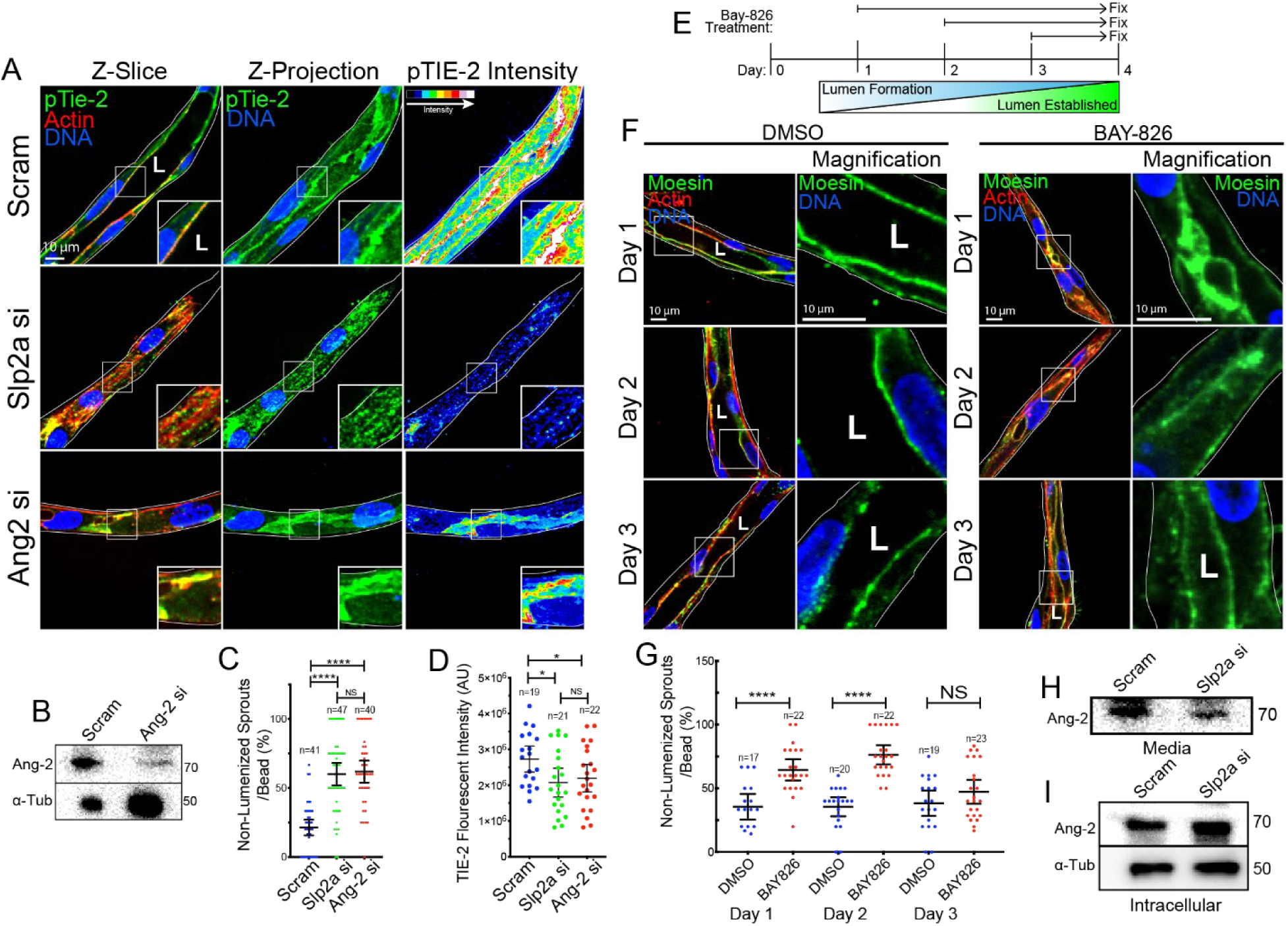
Slp2a mediates Ang-2 secretion and downstream Tie-2 signaling during lumen formation. **A,** Images of scramble (scram), Slp2a and Ang-2 siRNA (si) knockdown sprouts stained with actin and phosphorylated Tie-2 (pTie-2). Last column pseudo-colored to visualize antibody staining intensity. **B,** Representative confirmation of Ang-2 knockdown via western blot probed for Ang-2 and alpha-tubulin (α-Tub). n=3. **C**, Quantification of percent non-lumenized sprouts between indicated groups. **D,** Quantification of pTie-2 fluorescent intensity between indicated groups. **E,** Schematic of experimental setup with Tie-2 inhibitor Bay-826 to determine effect of Tie-2 signaling on lumen formation. **F**, Representative images of sprouts treated with Bay-826 or DMSO (control) on indicated day and stained for luminal marker moesin and actin. **G**, Quantification of percent non-lumenized sprouts between indicated groups. **H**, Representative western blotting for Ang-2 secretion into culture media between indicated conditions. n=3. **I,** Representative western blot probing for intracellular Ang-2 by indicated groups. n=3. All experiments use human umbilical vein endothelial cells. AU= arbitrary unit. In all panels L denotes lumen; white box denotes magnification; white lines denotes exterior of sprout; NS= not significant, values are means +/- SEM; n= individual sprouts across three experimental repeats; significance: *P<0.05, ****P<0.00005, NS=Not Significant. Statistical significance was assessed with a 1-way ANOVA followed by a Dunnett multiple comparisons test.

### Slp2a/b and Tie-2 signaling are required for lumen formation in developing zebrafish blood vessels

To confirm our results *in vivo* we turned to the zebrafish model of vascular development. Zebrafish blood vessel development is an established model of vascular lumen formation demonstrating stereotyped blood vessel morphology with an easily identifiable lumen cavity [33]. Furthermore, zebrafish are exceptionally well-suited for gene knockout studies using CRISPR/Cas9 editing [34-37]. Using CRISPR/Cas9 targeting as previously described [34] we knocked out both paralogs of Slp2 (A and B). Our sequence analysis showed an 75% indel formation with a ∼50% reduction in both Slp2a and Slp2b transcripts (**Fig. 7A**). Knockouts, singly or in combination, did not alter larvae body plan or growth kinetics (**Fig. S8A**). Inspection of the intersomitic vessels (ISVs) showed an increase in non-lumenzed ISVs particularly in the Slp2a/b crispants compared with scrambled sgRNA injected controls (**Fig. S8B**). Interestingly, ISVs were fully formed, connecting to the dorsal longitudinal anastomotic vessels (DLAV) at 36 hours post fertilization (hpf) (**Fig. S8B**), suggesting migratory processes were unaffected. Also, we did not observe any major differences in survival, ectopic or incomplete ISVs at 36 or 48 hpf (**Fig. S8C- H**). Next, we used microangiography to demarcate blood vessels with an open, contiguous luminal cavity. This method allowed us to conclusively assess whether vascular lumens were open, narrowed or non-existent (**Fig. 7B**). Compared with controls, knockout of both Slp2 paralogs significantly increased the percentage of non-perfused ISVs (Fig. 7C,D). This result tracked with significantly elevated number of non-lumenized ISVs in the double Slp2a/b KO compared with individual paralogs KOs or controls (**Fig. 7E**). This increase in non-lumenized vessels was independent of defects in ISVs as there was no difference in the number of formed ISVs between groups (**Fig. 7F**). Using a different approach, we marked the apical membrane of the forming ISVs by expressing PHluorin-podocalyxin. PHluorin is a GFP variant that is non-fluorescent in acidified vesicles, but fluorescence is rescued at neutral pH following membrane fusion [38]. This method allowed us to visualize podocalyxin that was inserted into the plasma membrane, clearing defining the apical surface (**Fig. S9A**). Using this marker, knockout of Slp2a/b also demonstrated a defined collapse of the apical membrane marked by a loss of PHluorin-podocalyxin signal (**Fig. S9B**). Overall, this data suggests that Slp2a/b is necessary for lumen formation *in vivo*.

**Figure 7.**
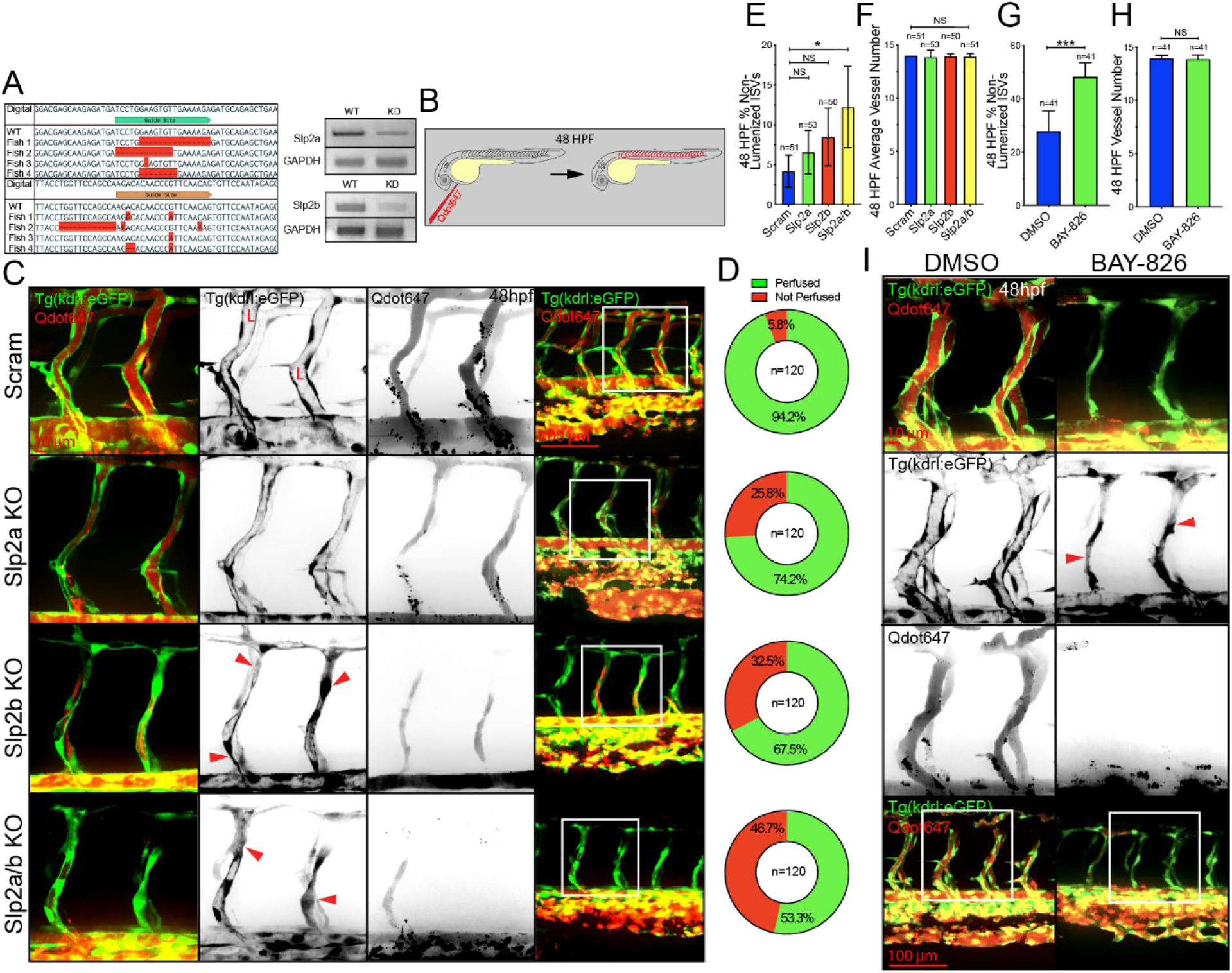
Genetic knockdown of Slp2a/b in zebrafish blunts lumen formation. **A,** Sequence of fish targeting sites after CRISPR/Cas9 gene editing aligned to wild-type (WT) sequence. Bands are RT-PCR analysis of mRNA transcript levels in zebrafish injected with indicated sgRNA normalized to a GAPDH expression control. 4 fish per guide were analyzed for targeting analysis and 20 fish per condition were homogenized for RT-PCR over three experimental repeats. In a pooled sample expression of Slp2a/b KD is ∼50% reduced relative to WT. **B,** Cartoon schematic of microangiography. Zebrafish were perfused with Quantum dot 647 (Qdot647) to highlight the vascular lumen cavity. **C,** 48 hours post fertilization (hpf) zebrafish tg(kdrl:GFP) perfused with Qdot 647 between indicated conditions. Arrowheads indicate sites of lumen collapse. **D,** Quantification of percentage of perfused intersomitic vessels (ISVs) between indicated crispant groups. n=numbers of ISVs over three experimental replicates. **E,** Quantification of percentage of non-lumenized ISVs at 48 hpf. N-value represents individual fish. **F,** Quantification of number of ISVs between indicated crispant groups. N-value represents individual fish. **G**, Quantification of non-lumenized ISVs between DMSO (control) and Bay-826 (small molecule Tie-2 inhibitor). N- values represent individual fish over three experimental repeats. **H**, Quantification of ISV number between indicated groups. N-values represent individual fish over three experimental repeats. **I**, Representative images of 48 hpf zebrafish ISVs perfused with Qdot 647 between indicated conditions. In all panels L denotes lumen; arrowheads denote lumen failure; values are means +/- SEM; significance: *P<0.05, ***P<0.001, NS= not significant. Statistical significance was assessed with an unpaired students t-test or 1-way ANOVA followed by a Dunnett multiple comparisons test.

To test if Slp2a or Rab27a maintained their cellular localization patterns in forming zebrafish blood vessels, we mosaically overexpressed tagged-version of Rab27a, pro-vWF [24] and Slp2a. Similar to our *in vitro* results, we observed both vWF and Rab27a were located in intracellular puncta in zebrafish ISVs (**Fig. S9C**). Interestingly, overexpression of pro-vWF also accumulated in the luminal space, suggestive of clot formation. Overexpression of tagged-Slp2a also demonstrated similar localization patterns as compared with our *in vitro* sprouting model. Both Slp2a WT and Slp2a-C2AB were membranous, while the Slp2a-ΔC2AB mutant was cytoplasmic (**Fig. S9D**).

To further substantiate our *in vitro* results, we applied the Tie-2 inhibitor Bay-826 used previously to larvae at 24 hpf, and then quantified for lumen defects at 48 hpf. During this time ISVs sprout dorsally from the dorsa aorta and lumenize prior to forming the DLAV. In line with our *in vitro* results, we observed a significant increase in the number of non-lumenized ISVs compared to a vehicle control (Fig. 7G, I). Again, these results were independent of alterations in ISV number or body plan (**Fig. 7H; S8I**). Cumulatively, our data indicates that Tie-2 signaling is required for blood vessel lumenogenesis *in vivo* (**Fig. 8**).

**Figure 8.**
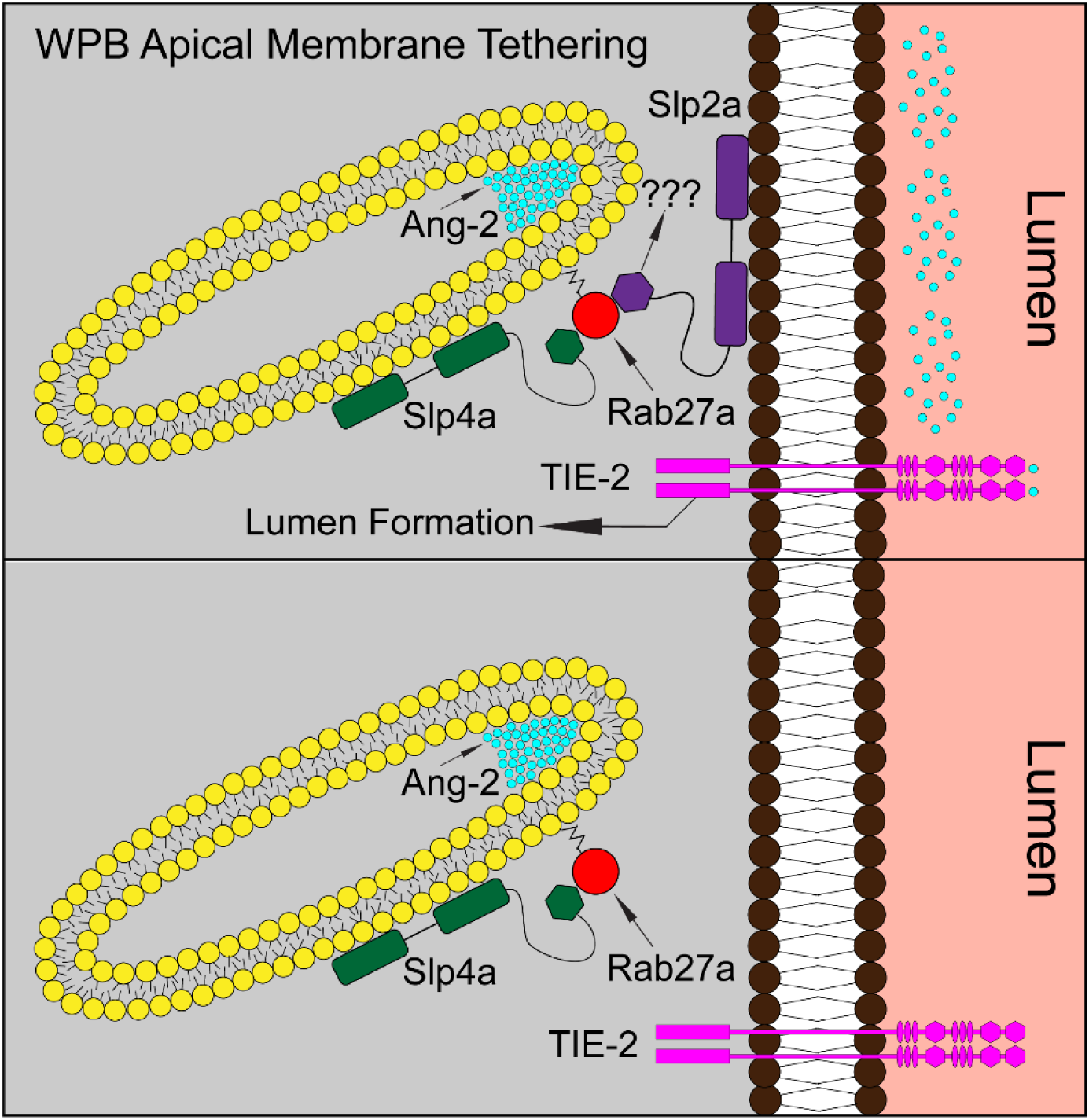
Proposed model of Slp2a function in vascular lumen formation. Top panel cartoon representation of Slp2a acting as a tether at the apical membrane for WPBs binding Rab27a. Ang-2 housed within the WPBs are then successfully targeted to the apical membrane and secreted. Once secreted, Ang-2 binds the receptor Tie-2 leading to downstream signaling promoting lumen formation and sprout stabilization. The bottom panel cartoon is lacking Slp2a and therefor WPBs fail to properly exocytose cargo, reducing Ang-2 autocrine signaling and Tie- 2 signaling cascade.

## DISCUSSION

In the present investigation we demonstrate that Slp2a is required for vascular lumen formation. Our results establish that Slp2a resides at the apical membrane where it can tether Rab27a positive vesicles. Unique to endothelial tissue, Slp2a controls the fusion of WPBs for secretion of their contents into the luminal space during angiogenesis. Ang-2 is a Tie-2 receptor ligand that is selectively exocytosed from WPB secretory granules and is necessary for proper blood vessel development in specific contexts [39]. In the absence of Slp2a, WPB contents cannot fuse with the apical membrane precluding the release of Ang-2, diminishing Tie-2 signaling necessary for proper lumen formation (**Fig. 8**). Overall, our results demonstrate that Slp2a is required for targeting secretory vesicles to the apical membrane during vascular lumen development and a core component of the WPB secretory pathway.

A fundamental morphogenic program during blood vessel development is the creation of a continuous lumen as a conduit for blood flow [6, 7, 35]. This feat requires the establishment of apicobasal membrane polarities via the recruitment of lipids and proteins to differential plasma membrane domains. In the present investigation we demonstrate that Slp2a is necessary for lumen formation in vascular development by regulating the secretion of WPB vesicles at the apical membrane. We believe as all blood vessels form a lumen during their development, Slp2a could be a major contributor to lumenogenesis in all vessel beds ranging from small capillaries to large arteries. Interestingly, and unlike epithelial tissue, Slp2a’s trafficking partner Rab27a does not deliver podocalyxin in ECs, but is intimately involved in WPB granule secretion. This novel association highlights an endothelial-specific function as ECs are the predominant harbor of WPBs due to their central role in hemostasis. This finding brings into question, if Rab27a is not transporting podocalyxin, what other(s) trafficking pathway has been evoked in its place?

Rab27a is one of the best characterized exocytic Rab GTPase family members [14, 36, 37]. Rab27a function in vascular tissue has been shown to regulate VEGFR1 trafficking by directing palmitoylation, controlling receptor recycling or through receptor degradation [19]. In the current investigation, Rab27a localized only to WPBs – we did not detect any obvious receptor labeling. Our findings were congruent with Nightingale et al. [18] in which Rab27a was a negative regulator of WPB exocytosis. Our results show that when Rab27a is ablated, WPB cargo is readily released; however, in the absence of Slp2a, Rab27a-evoked exocytosis is halted at the apical membrane. If Rab27a binding to Slp2a was a requisite for WPB cargo secretion, then knockdown of Rab27a would have also blunted vWF and Ang-2 release, we did not observe this. Thus, the exact mechanism by which Slp2a is tethering WPBs at the apical membrane remains unknown, we can only conclude that it is required for this process. To this effect, Slp2a could be also required for membrane fusion, such as in SNARE [38], Unc13 [40], VAMP [41] mediated events and not simply involved in tethering WPBs adjacent to the apical membrane. Additionally, other Ca^2+^-dependent synaptotagmins could be aiding in exocytic events at the luminal membrane [42]. Indeed, further experiments are required to fully understand Slp2a’s role in WPB docking and fusion at the apical membrane as well as its other potential functions as an exocytic regulator during angiogenic development.

The spatiotemporal regulation of WPB trafficking is complex demonstrating both constitutive and induced exocytic behaviors in ECs. In addition to this complexity, the contents of WPBs are vast, with as many as 40 proteins cited as WPB cargo [43]. It has been reported that subpopulations of WPB granules exhibit preferential housing of certain proteins, to the exclusion of others [28]. For example, P-selectin and Ang-2 occupy mutually exclusive WPB populations [44]. Our results support this notion as we observed some selectivity in Ang-2 localization compared with vWF-positive WPBs. These observations necessitate the notion of unique trafficking programs that WPBs execute to complement their growing role as a dynamic exocytic depot. We also observed that the non-membrane binding Slp2a-ΔC2AB mutant decorated virtually all vWF and Ang-2 positive vesicles, indicating that regardless of the WPB sub- population, Slp2a is likely a requirement for apical membrane tethering and secretion.

Due to the continuously growing body of new data, Ang-2’s exact contribution to angiogenesis has yet to be resolved and is at present considered context dependent. Both Ang- 1 and Ang-2 can bind the Tie-2 receptor. Initial Ang-2 investigations provided evidence for Ang-2 solely functioning as an antagonist of Ang-1, in which Ang-1 binding promoted Akt signaling, fortifying vascular barrier function [30, 44, 45]. Contrary to these results, later studies showed, in a pathophysiological context, Ang-2 expression can promote Tie-2 activation as well as interacting with integrins [46]. This phenomenon has been observed during tumor-induced angiogenesis as well as in inflamed endothelium [32, 47]. Further complicating this interplay is the unique expression patterns between ECs and pericytes. Ang-1 is primarily secreted by pericytes whereas Ang-2 is released from the endothelium [48, 49]. In the absence of pericyte-derived Ang-1, we observed that knockdown of Ang-2 drastically affected vascular lumen formation. Furthermore, reduction of Ang-2 resulted in lower phosphorylated Tie-2 levels at the apical membrane in vascular sprouts, signifying an activating role in our model. Given these results, it is tempting to speculate that our fibrin-bead sprouting model may be more akin to tumor angiogenesis in which Ang-2 is converted to a Tie-2 activating ligand in the absence of pericyte-derived Ang-1. Our experiments also demonstrate elevated Ang-2 exocytosis into the luminal space during the beginning phases of lumen development and a reduction of luminal Ang-2 after sprouts were lumenized. To our knowledge, this is the first direct evidence of a graded secretion of Ang-2 into the lumen space, supportive of a requirement for Ang-2-mediated Tie-2 activation during lumen formation. Additionally, we provide new evidence for Slp2a acting as a gatekeeper for exocytic events involving WPB fusion at the apical membrane.

In summary, the present study provides a novel characterization of Slp2a and its role as an upstream apical membrane tether required for vascular lumen formation. Our evidence highlights a direct association between Slp2a, Rab27a and WPBs in which Slp2a functions to facilitate WPB secretion. Our results also show that WPB-housed Ang-2 is a critical secreted factor for physiological progression of vascular lumen formation. Cumulatively, Slp2a is a major upstream apical membrane protein controlling regulated secretion programs during angiogenic development, which may have other uncharacterized roles in both angiogenesis and adult blood vessel homeostasis controlling trafficking at the apical membrane.

## ACKNOWLEDGMENTS

We would like to thank Dr. J.T. Blankenship for critical reading of the manuscript.

## CONTRIBUTIONS

C.R.F, S.C and E.J.K performed all experiments. C.R.F and E.J.K wrote the manuscript.

## SUPPLEMENTAL MATERIALS AND METHODS

### Zebrafish Maintenance and Stock

Zebrafish were raised and maintained in accordance with institutional guidelines set by The University of Denver and national animal welfare regulations. Zebrafish experiments were approved by the University of Denver’s IACUC committee.

### Zebrafish Transgenics

The transgenic lines used in this study include Tg(kdrl:GFP) [1] and Tg(kdrl:mCherry)[2]. Tol2- mediated transgenesis was used to generate mosaic intersomitic blood vessels as previously described [3, 4]. Briefly, Tol2 transposase mRNA were synthesized (pT3TS-Tol2 was a gift from Stephen Ekker, Addgene plasmid # 31831) [5] using an SP6 RNA polymerase (mMessage mMachine, ThermoFisher). A total of 400ng of transposase and 200ng of plasmid vector were combined and brought up to 10μL with phenol red in ddH_2_O. The mixture was injected into embryos at the 1-2 cell stage. Injected zebrafish were screened for mosaic expression at 48 hpf and imaged. CRISPR/cas9-mediated knockouts were performed as previously described [6]. Briefly, equal volumes of chemically synthesized AltR® crRNA (100 μM) and tracrRNAr RNA (100 μM) were annealed by heating and gradual cooling to room temperature. Thereafter the 50:50 crRNA:tracrRNA duplex stock solution was further diluted to 25 μM using supplied duplex buffer. Prior to injection 25 μM crRNA:tracrRNA duplex stock solution was mixed with 25 μM Cas9 protein (Alt-R® S.p. Cas9 nuclease, v.3, IDT) stock solution in 20mM HEPES-NaOH (pH 7.5), 350mM KCl, 20% glycerol) and diluted to 5μM by diluting with water. Prior to microinjection, the RNP complex solution was incubated at 37°C, 5 min and then placed on ice. The injection mixture was micro-injected into 1-2 cell stage embryos. Confirmation of CRISPR-mediated gene knockout was validated by RT-PCR. Approximately 20 embryos at 48 hpf were homogenized and dissolved in TRIZOL reagent (Sigma Aldrich) according to the manufacturer’s instructions. Reverse transcription was achieved by using the Biosystems High-Capacity cDNA Reverse Transcription Kit (ThermoFisher) and then PCR was used with relevant controls (See table 1.1 for DNA oligos and sgRNA sequences used) to determine transcript levels. Crispant DNA was retrieved via PCR and subjected to sanger sequencing to visualize indel formation.

### Zebrafish Microangiography

Embryos were (anesthetized) with 1% tricaine for approximately 20 minutes prior to perfusion. Embryos were then loaded ventral side up onto an injection agarose facing the injection needle. Prior to injection Qdots (ThermoFisher) were prepped by 20 minutes of sonication. Qdots were loaded into a pulled capillary needle connected to an Eppendorf CellTram and 1-3μl of perfusion solution was injected into the pericardial cavity. Once successfully perfused, embryos were embedded in 0.7% low melt agarose and imaged promptly. Images were taken on a Nikon Eclipse Ti inverted microscope equipped with a CSU-X1 Yokogawa spinning disk field scanning confocal system and a Hamamatusu EM-CCD digital camera using either Nikon Apo LWD 20x NA 0.95 or Nikon Apo LWD 40x NA 1.15 water objective.

### Zebrafish Live Imaging and Quantification

All zebrafish presented were imaged at 36 and 48hpf. Prior to imaging, embryos were treated with 1% Tricaine for 20 minutes and afterwards embedded in 0.7% low melt agarose. Live imaging of Zebrafish intersomic vessels (ISVs) was performed using a spinning-disk confocal microscopy system mentioned above. Zebrafish embryos were quantified at 36hpf and 48hpf. ISVs that were analyzed were between the end of the yolk extension and tail. Parameters measured included ISV number, number non-lumenized vessels (no visible separation between opposing endothelial cells in vessels), number of ectopic vessels (extra vessels), and number of incomplete vessels (vessels that do not contact the dorsal longitudinal anastomotic vessel at 48 hpf).

### Lentivirus and Adenovirus Generation and Transduction

Lentivirus was generated by using the LR Gateway Cloning method [7]. Genes of interest and fluorescent proteins were isolated and incorporated into a pME backbone via Gibson reaction [8]. Following confirmation of the plasmid by sequencing the pME entry plasmid was mixed with the destination vector and LR Clonase. The destination vector used in this study was pLenti CMV Neo DEST (705-1) (gift from Eric Campeau & Paul Kaufman; Addgene plasmid #17392). Once validated, the destination plasmids were transfected with the three required viral protein plasmids: pMDLg/pRRE (gift from Didier Trono; Addgene plasmid # 12251), pVSVG (gift from Bob Weinberg; Addgene plasmid #8454) and psPAX2 (gift from Didier Trono; Addgene plasmid #12260) into HEK 293 cells. The transfected HEKs had media changed 4 hours post transfection. Transfected cells incubated for 3 days and virus was harvested.

Adenoviral constructs and viral particles were created using the Adeasy viral cloning protocol [9]. Briefly, transgenes were cloned into a pShuttle-CMV plasmid (gift from Bert Vogelstein; Addgene plasmid #16403) via Gibson Assembly. PShuttle-CMV plasmids were then digested overnight with MssI (ThermoFisher) and Linearized pShuttle-CMV plasmids were transformed into the final viral backbone using electrocompetent AdEasier-1 cells (gift from Bert Vogelstein; Addgene, #16399). Successful incorporation of pShuttle-CMV construct into AdEasier-1 cells confirmed via digestion with PacI (ThermoFisher). 5000 ng plasmid was then digested at 37°C overnight, then 85°C for 10 minutes and transfected in a 3:1 polyethylenimine (PEI, Sigma):DNA ratio into 70% confluent HEK 293A cells (ThermoFisher) in a T-25 flask.

Over the course of 2-4 weeks, fluorescent cells became swollen and budded off the plate. Once approximately 70% of the cells had lifted off the plate, cells were scraped off and spun down at 2000 rpm for 5 minutes in a 15 mL conical tube. The supernatant was aspirated, and cells were resuspended in 1 mL PBS. Cells were then lysed by 3 consecutive quick freeze-thaw cycles in liquid nitrogen, spun down for 5 minutes at 2000 rpm, and supernatant was added to 2qty 70% confluent T-75 flasks. Propagation continued and collection repeated for infection of 10-15cm dishes. After collection and 4 freeze thaw cycles of virus collected from 10-15cm dishes, 8 mL viral supernatant was collected and combined with 4.4 g CsCl (Sigma) in 10 mL PBS. Solution was overlaid with mineral oil and spun at 32,000 rpm at 10°C for 18 hours. Viral fraction was collected with a syringe and stored in a 1:1 ratio with a storage buffer containing 10 mM Tris, pH 8.0, 100 mM NaCl, 0.1 percent BSA, and 50% glycerol. HUVEC were treated with virus for 16 hours at a 1/1000 final dilution in all cell culture experiments.

### Immunoblotting & Protein Pull-Down

HUVEC culture was trypsinized and lysed using Ripa buffer (20 mM Tris-HCl [pH 7.5], 150 mM NaCl, 1 mM Na_2_EDTA, 1 mM EGTA, 1% NP-40, 1% sodium deoxycholate, 2.5 mM sodium pyrophosphate, 1 mM β-glycerophosphate, 1 mM Na_3_VO_4_, 1 µg/mL leupeptin) containing 1x ProBlock™ Protease Inhibitor Cocktail -50 (GoldBio). Total concentration of protein in lysate was quantified using the Pierce™ BCA Protein Assay Kit measured at 562 nm and compared to a standard curve. 20-50 µg protein was prepared in 0.52 M SDS, 1.2 mM bromothymol blue, 58.6% glycerol, 75 mM Tris pH 6.8, and 0.17 M DTT. Samples were boiled for 10 minutes, then loaded in a 7-12% SDS gel and run at 150 V. Protein was then transferred to Immun-Blot PVDF Membrane (BioRad) at 4°C, 100 V for 1 hour 10 minutes. Blots were blocked in 2% milk proteins for 1 hour, then put in primary antibody at specified concentrations overnight. After 3 10-minute washes with PBS, secondary antibodies at specified concentrations were applied for 4 hours. After 3 additional PBS washes, blots were developed with ProSignal® Pico ECL Spray.

For Rab27a pull-down experiments, GST-Slp2a was grown overnight in 50 mL of Laria- Bertani broth in NiCo21 E Coli (NEB). The following day the overnight culture was transferred to 1L of terrific buffer. The culture was monitored for growth and induced at OD600 with IPTG (GoldBio, 12481) at a final concentration of 100uM. Following induction, bacteria were incubated for an additional 3 hours. Induced cells were collected and pelleted, with the pellet resuspended in cold PBS containing 1mg/ml lysozyme and 1x ProBlock™ Protease Inhibitor Cocktail -50 and then sonicated to lyse bacteria. Cell lysate was clarified by centrifugation and glutathione agarose resin (GoldBio) was added to affinity purify the GST-Slp2a. After incubation, agarose resin was washed 2-3 times with PBS and stored at -20°C.

## MAJOR RESOURCE TABLE

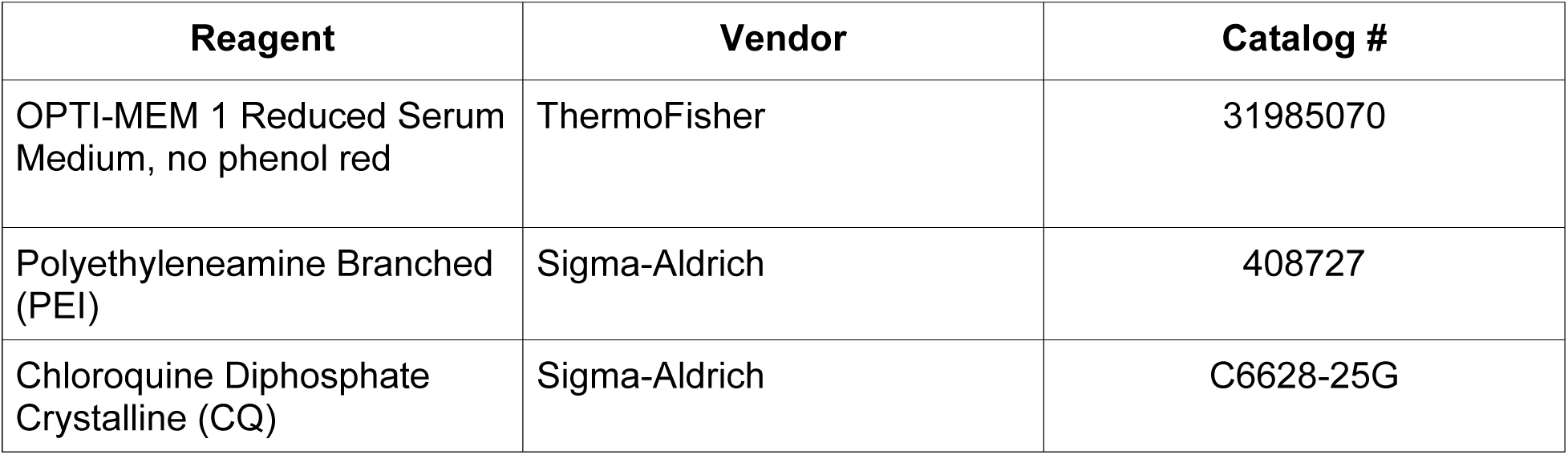
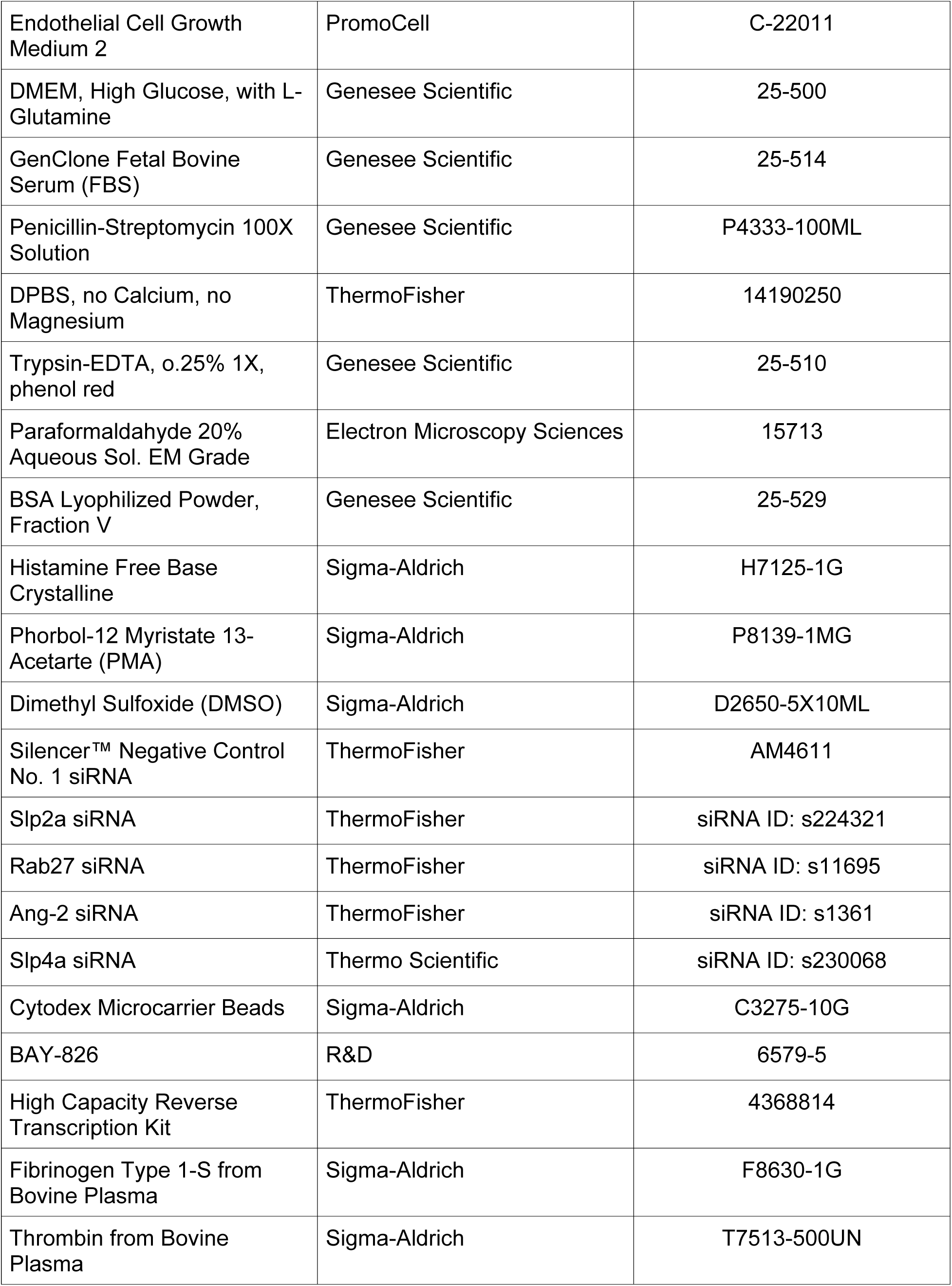
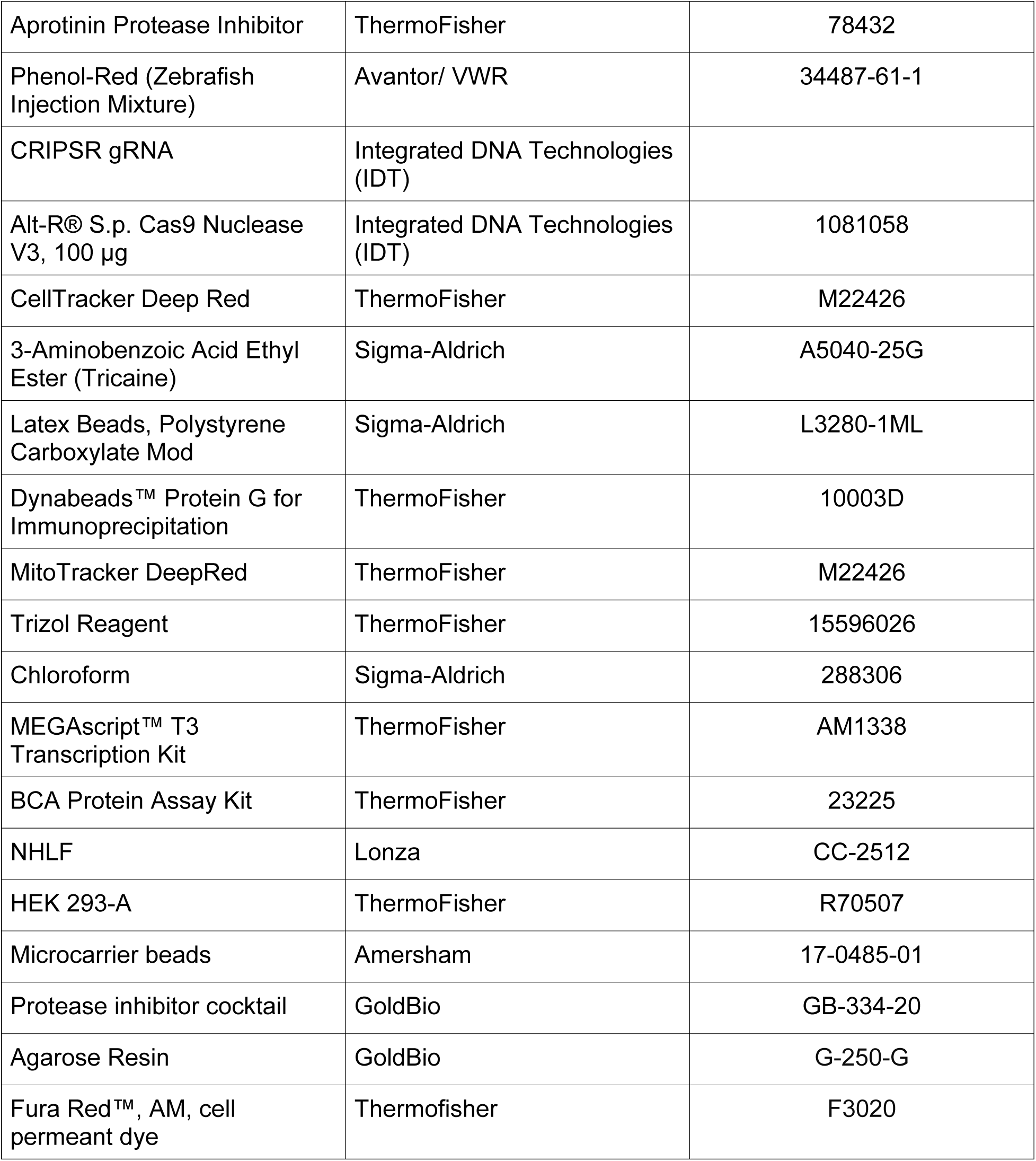

| Reagent | Vendor | Catalog # |
| --- | --- | --- |
| OPTI-MEM 1 Reduced Serum Medium, no phenol red | ThermoFisher | 31985070 |
| Polyethyleneamine Branched (PEI) | Sigma-Aldrich | 408727 |
| Chloroquine Diphosphate Crystalline (CQ) | Sigma-Aldrich | C6628-25G |
| Endothelial Cell Growth Medium 2 | PromoCell | C-22011 |
| DMEM, High Glucose, with L- Glutamine | Genesee Scientific | 25-500 |
| GenClone Fetal Bovine Serum (FBS) | Genesee Scientific | 25-514 |
| Penicillin-Streptomycin 100X Solution | Genesee Scientific | P4333-100ML |
| DPBS, no Calcium, no Magnesium | ThermoFisher | 14190250 |
| Trypsin-EDTA, 0.25% 1X, phenol red | Genesee Scientific | 25-510 |
| Paraformaldehyde 20% Aqueous Sol. EM Grade | Electron Microscopy Sciences | 15713 |
| BSA Lyophilized Powder, Fraction V | Genesee Scientific | 25-529 |
| Histamine Free Base Crystalline | Sigma-Aldrich | H7125-1G |
| Phorbol-12 Myristate 13-Acetate (PMA) | Sigma-Aldrich | P8139-1MG |
| Dimethyl Sulfoxide (DMSO) | Sigma-Aldrich | D2650-5X10ML |
| Silencer™ Negative Control No. 1 siRNA | ThermoFisher | AM4611 |
| Slp2a siRNA | ThermoFisher | siRNA ID: s224321 |
| Rab27 siRNA | ThermoFisher | siRNA ID: s11695 |
| Ang-2 siRNA | ThermoFisher | siRNA ID: s1361 |
| Slp4a siRNA | Thermo Scientific | siRNA ID: s230068 |
| Cytodex Microcarrier Beads | Sigma-Aldrich | C3275-10G |
| BAY-826 | R&D | 6579-5 |
| High Capacity Reverse Transcription Kit | ThermoFisher | 4368814 |
| Fibrinogen Type 1-S from Bovine Plasma | Sigma-Aldrich | F8630-1G |
| Thrombin from Bovine Plasma | Sigma-Aldrich | T7513-500UN |
| Aprotinin Protease Inhibitor | ThermoFisher | 78432 |
| Phenol-Red (Zebrafish Injection Mixture) | Avantor/ VWR | 34487-61-1 |
| CRIPSR gRNA | Integrated DNA Technologies (IDT) |  |
| Alt-R® S.p. Cas9 Nuclease V3, 100 µg | Integrated DNA Technologies (IDT) | 1081058 |
| CellTracker Deep Red | ThermoFisher | M22426 |
| 3-Aminobenzoic Acid Ethyl Ester (Tricaine) | Sigma-Aldrich | A5040-25G |
| Latex Beads, Polystyrene Carboxylate Mod | Sigma-Aldrich | L3280-1ML |
| Dynabeads™ Protein G for Immunoprecipitation | ThermoFisher | 10003D |
| MitoTracker DeepRed | ThermoFisher | M22426 |
| Trizol Reagent | ThermoFisher | 15596026 |
| Chloroform | Sigma-Aldrich | 288306 |
| MEGAscript™ T3 Transcription Kit | ThermoFisher | AM1338 |
| BCA Protein Assay Kit | ThermoFisher | 23225 |
| NHLF | Lonza | CC-2512 |
| HEK 293-A | ThermoFisher | R70507 |
| Microcarrier beads | Amersham | 17-0485-01 |
| Protease inhibitor cocktail | GoldBio | GB-334-20 |
| Agarose Resin | GoldBio | G-250-G |
| Fura Red™, AM, cell permeant dye | Thermofisher | F3020 |

## ANTIBODIES

| Target Antigen | Vendor or Source | Catalog No./ Clone | Working Concentration |
| --- | --- | --- | --- |
| Slp2a | ThermoFischer | PA524730 | 0.5ug/mL (1:1000) |
| Slp4a | ThermoFischer | PA551605 | 0.2 ug/mL<br>(1:1000) |
| Rab27a | Therm Fischer | PA551561 | 0.05ug/mL<br>(1:1000) |
| Ang2 | Thermo Fischer | PA129640 | 40mg/mL 1:500 |
| Alpha-tubulin | Abcam | ab52866 | 0.0648ug/mL<br>(1:10,000) |
| Moesin | Abcam | ab52490 | 0.05ug/mL<br>(1:1000) |
| VE-Cadherin | ThermoFischer | 14-1441-82 | 0.5ug/mL<br>(1:1000) |
| Podocalyxin | R&D | AF1658 | 15ug/mL (1:200) |
| Von Willebrand<br>Factor | Abcam | ab6994 | 10ug/mL<br>(1:1000) |
| Phosphorylated<br>TIE-2/TEK<br>(Tyr992) | Sigma Aldrich | ABF131 | 0.25 ug/mL<br>(1:1000) |
| Anti-HA-Tag,<br>Rabbit<br>Monoclonal | Sigma-Aldrich | SAB5600116-100UG | 5ug/mL |
| Alexa Fluor™<br>488 Phalloidin | ThermoFisher | A12379 | 1 uM (1:200) |
| Alexa Fluor™<br>647 Phalloidin | ThermoFisher | A22287 | 1 uM (1:200) |
| Alexa Fluor™<br>555 Phalloidin | ThermoFisher | A34055 | 1 uM (1:200) |
| Goat anti-Rabbit<br>IgG (H+L)<br>Secondary<br>Antibody, Alexa<br>Fluor 488 | ThermoFisher | A11008 | 1ug/mL (1:500) |
| Donkey anti-Rabbit IgG (H+L)<br>Secondary<br>Antibody, Alexa<br>Fluor 555 | ThermoFisher | A31572 | 1ug/mL (1:500) |
| Donkey anti-goat IgG (H+L) Secondary Antibody, Alexa Flour 488 | ThermoFisher | A11055 | 1ug/mL (1:500) |
| Donkey anti-Goat IgG (H+L) Cross-Adsorbed Secondary Antibody, Alexa Fluor 555 | ThermoFisher | A21432 | 1ug/mL (1:500) |
| Chicken anti-Rabbit IgG (H+L) Cross-Adsorbed Secondary Antibody, Alexa Fluor 647 | ThermoFisher | A21443 | 1ug/mL (1:500) |
| Goat Anti-Rabbit HRP | Genesee Scientific | 20-303 | 1ug/mL (1:500) |

## OLIGOS AND SGRNA

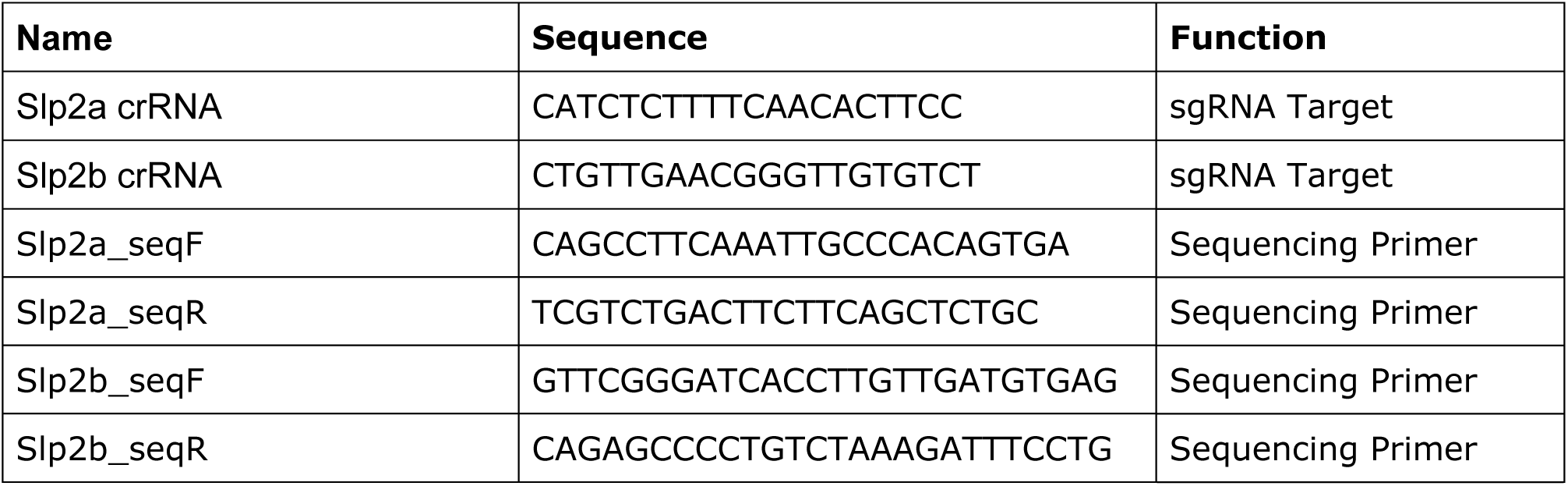

## SUPPLEMENTAL FIGURES

**Supplemental Figure 1.**
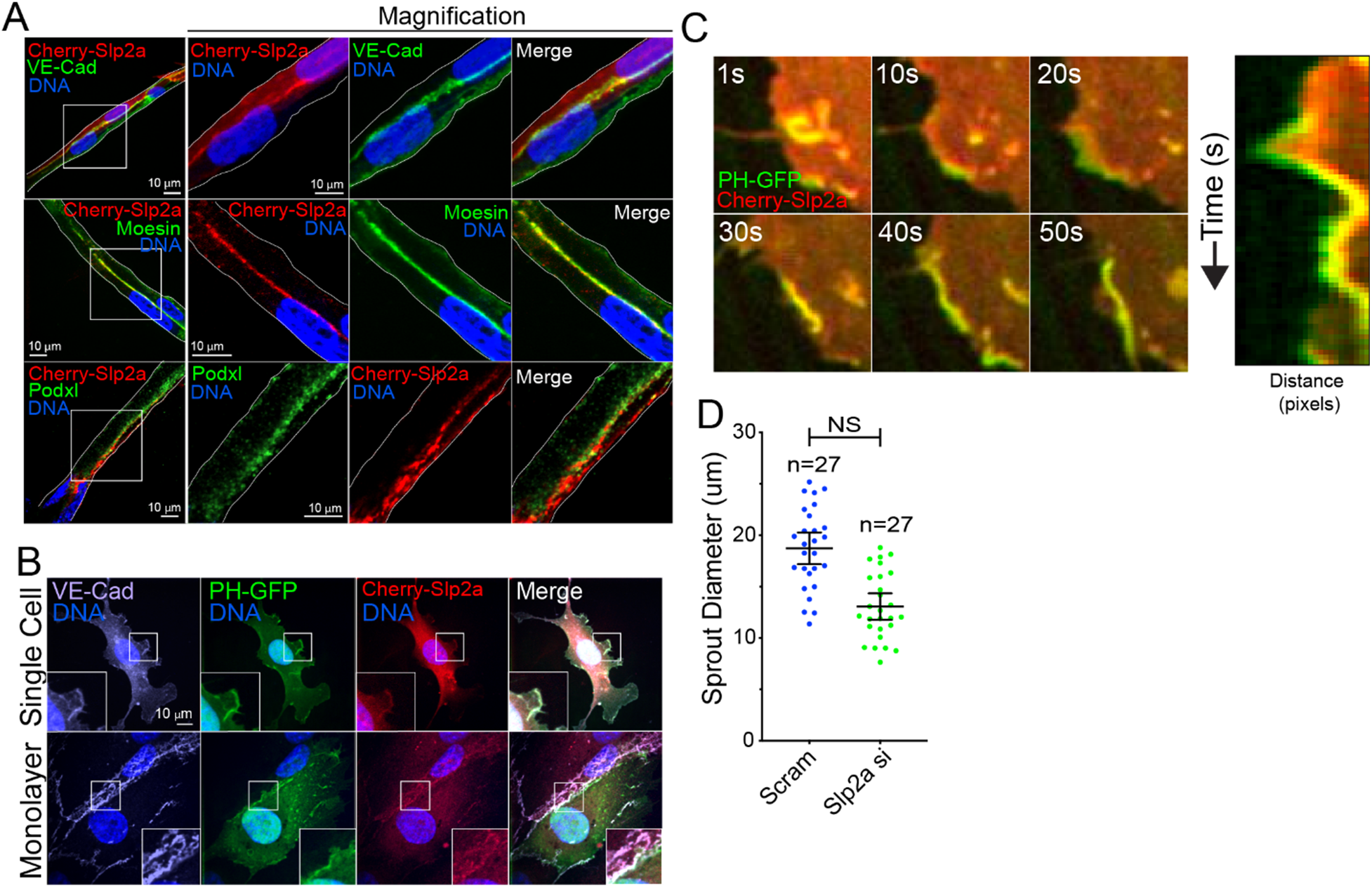
Slp2a localizes with PI(4,5)P_2_ at junctions. **A**, Sprouts expressing mCherry(Cherry)-tagged Slp2a fixed prior to lumen development and stained for VE-Cadherin (VE-cad), moesin, and podocalyxin (Podxl). **B**, Cells transduced with PH-GFP (PIP_2_ biosensor) and Cherry-Slp2a and stained as indicated. **C**, Live imaging of Cherry-Slp2a co-transduced with PH-GFP. Time points are shown in 10 second intervals with kymograph on right. **D,** Sprout diameter (um) of scramble (scram) and Slp2a siRNA(si) treated HUVECs. N-value represents individual sprouts over three experimental repeats. All experiments use human umbilical vein endothelial cells. In all panels L denotes lumen; white box denotes magnification; white lines denote exterior of sprout. NS= not significant, values are means +/- SEM; N= individual sprouts; significance: Statistical significance was assessed with unpaired t-test. All experiments were performed in triplicate.

**Supplemental Figure 2.**
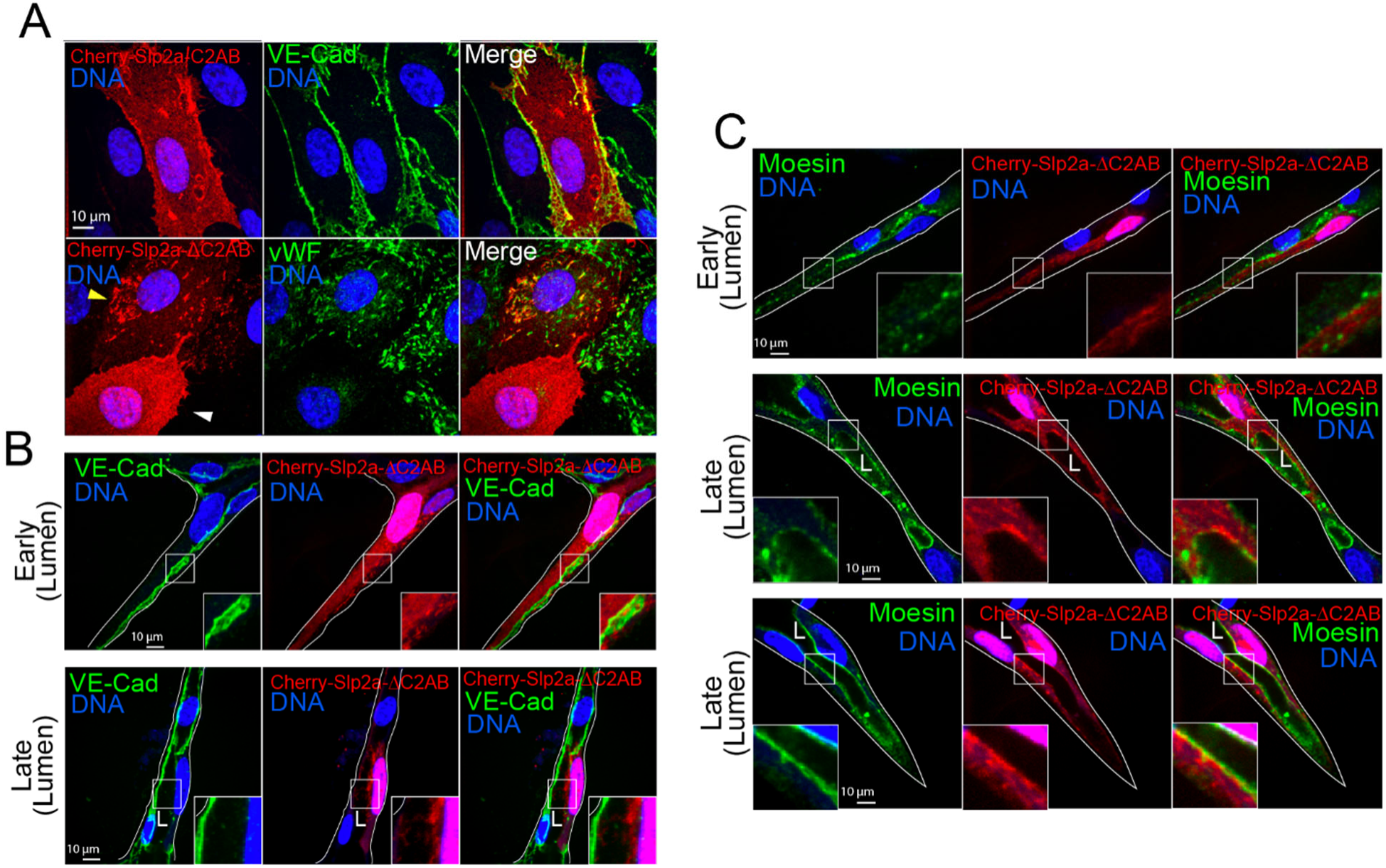
Slp2a-ΔC2AB Localization throughout lumen formation. **A,** Cells expressing mCherry(Cherry)-Slp2a-C2AB (top row) and stained for VE-Cadherin (VE-Cad). *Bottom row*, cells expressing Cherry-Slp2a-ΔC2AB and stained for von-Willebrand factor (vWF). Yellow arrow indicates cherry-Slp2a-ΔC2AB localization in cell expressing vWF and white arrow indicates cherry-Slp2a-ΔC2AB localization in cell lacking vWF expression. **B**, Cherry-Slp2a-ΔC2AB transduced sprouts fixed at early and late stages of lumen formation and stained for VE-Cad. **C**, Cherry-Slp2a-ΔC2AB expressing sprouts fixed at early and late stages of lumen formation. Sprouts were probed for moesin to highlight lumen cavity. White box denotes magnification. All experiments were performed in triplicate.

**Supplemental Figure 3.**
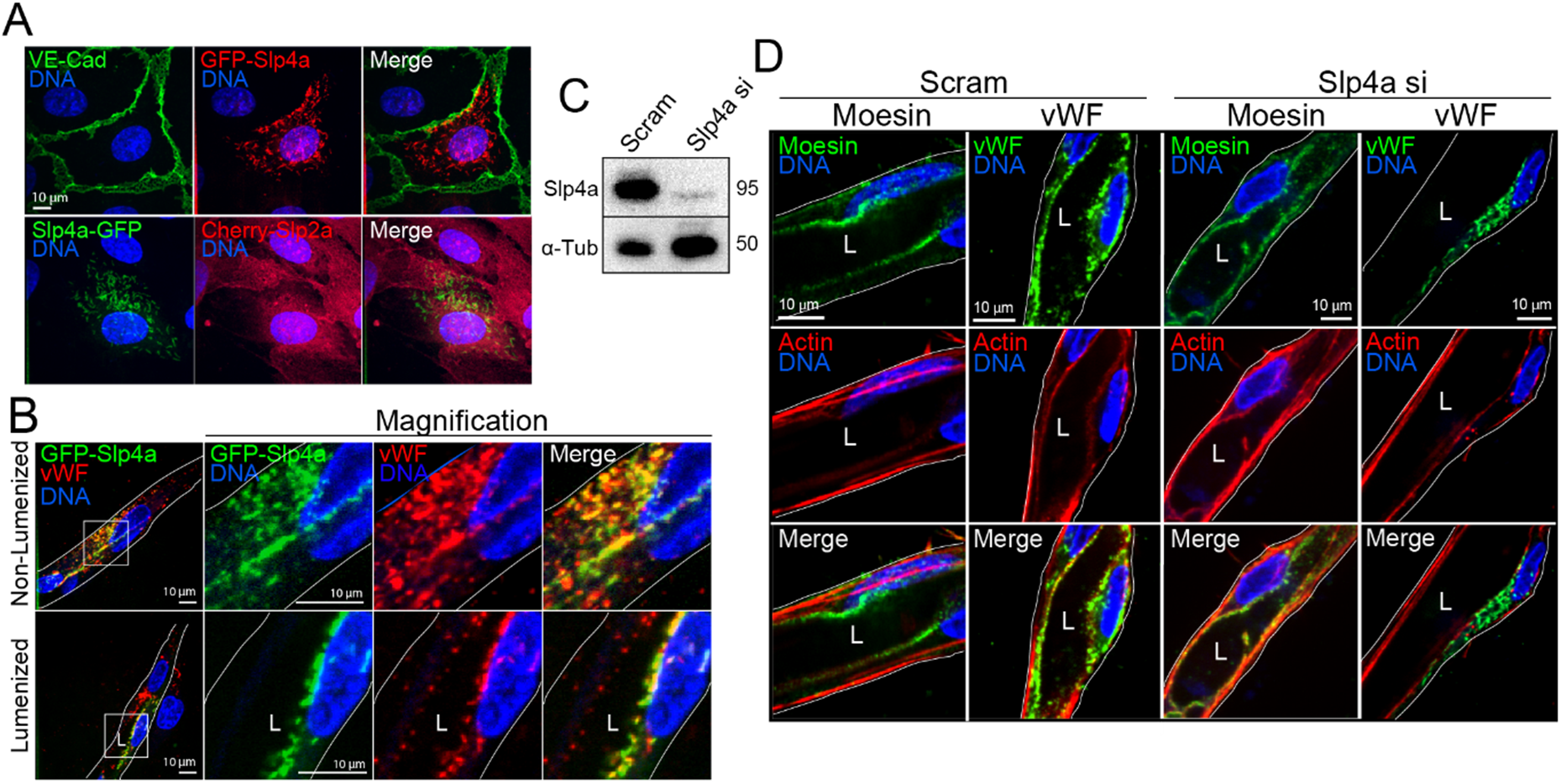
Slp2a, but not Slp4a, influences lumen development. **A**, Cells expressing GFP-Slp4a, mCherry(Cherr)-Slp2a, and stained for VE-Cadherin (VE-Cad). **B**, Sprouts expressing GFP-Slp4a and stained as indicated fixed before (non-lumenized) and after (lumenized) lumen formation. **C**, Representative western blot confirmation of Slp4a siRNA(si) knockdown. **D**, Images of Slp4a knockdown sprouts stained as indicated. All experiments use human umbilical vein endothelial cells. In all panels L denotes lumen; white box denotes magnification; white lines denote exterior of sprout. All experiments were performed in triplicate.

**Supplemental Figure 4.**
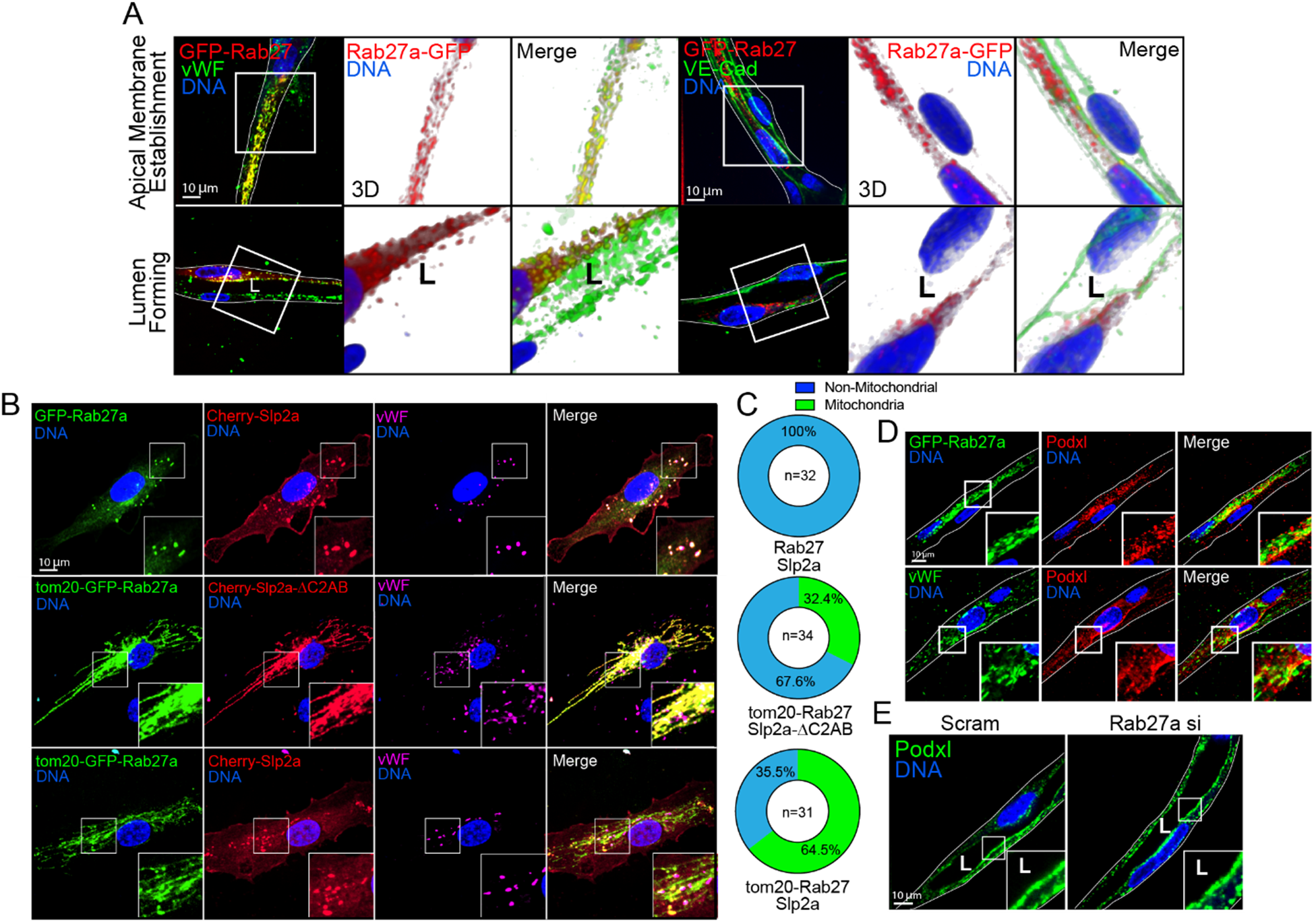
Rab27a localizes to WPBs and does not influence lumen formation. **A**, Sprouts expressing GFP-Rab27a and stained for VE-cadherin (VE-cad) and von Willebrand factor (vWF). Images were reconstructed in 3D. **B**, Three different scenarios for vWF localization endogenously and at the mitochondria. *Top row*, GFP-Rab27a co-expressed with Cherry-slp2a in cells stained for vWF. *Middle row*, tom20-GFP-Rab27a coexpressed with Cherry-Slp2a-ΔC2AB stained for vWF. *Bottom row*, tom20-GFP-Rab27a coexpressed with Cherry-Slp2a showing vWF colocalization at the mitochondria. **C**, Quantification of vWF localization at the mitochondria. N= individual sprouts. **D**, Localization of podocalyxin (Podxl) in sprouts relative to GFP-Rab27a and vWF. **E**, Scramble (scram) vs Rab27a siRNA (si)-treated sprouts stained for Podxl. All experiments use human umbilical vein endothelial cells. In all panels L denotes lumen; white box denotes magnification; white lines denote exterior of sprout. All experiments were performed in triplicate.

**Supplemental Figure 5.**
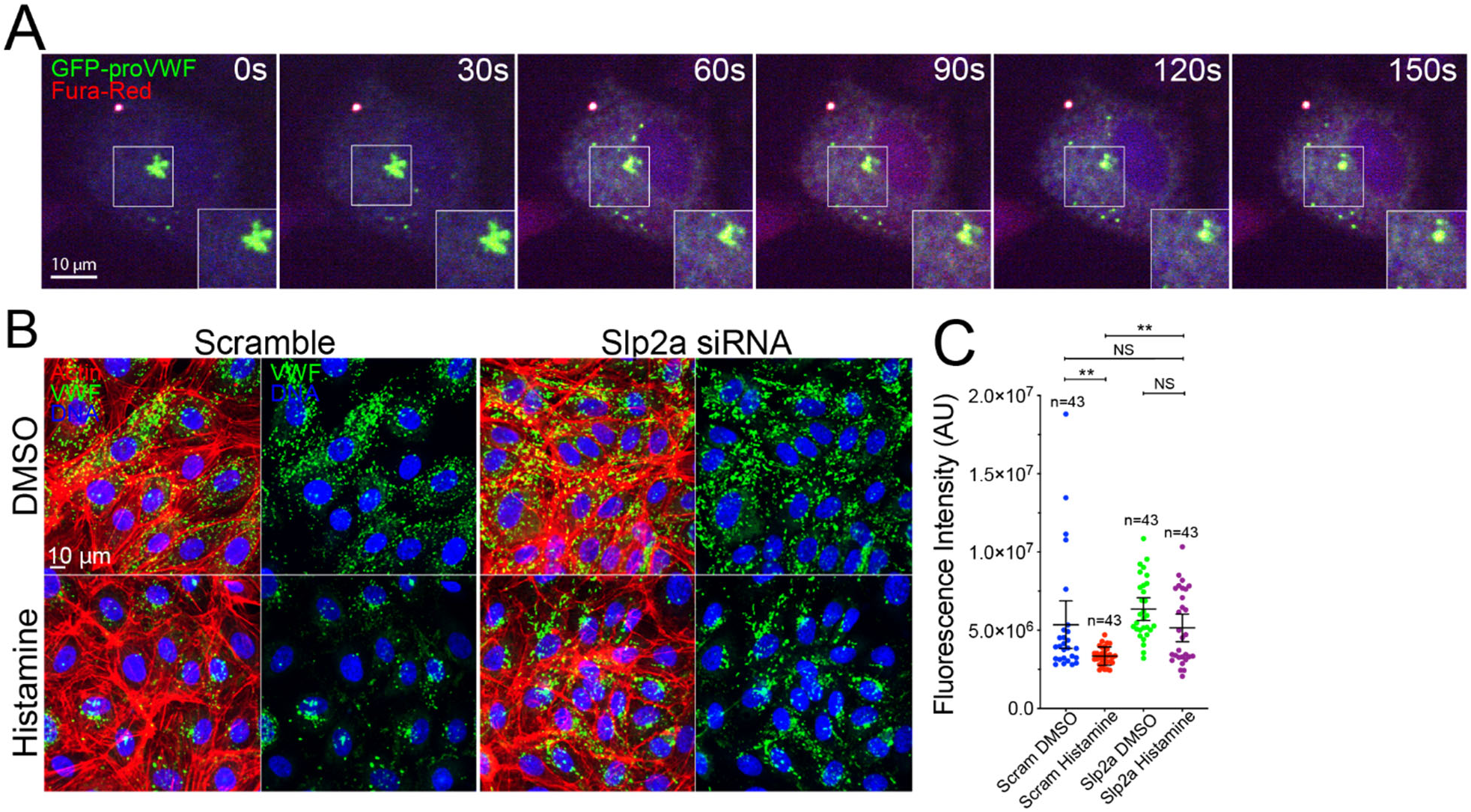
Slp2a is required for vWF secretion. **A,** Optimization experiment for histamine induced secretion. Fura-red indicator for calcium influx beginning at 0 seconds (histamine addition) to 150 seconds. Cell expressing GFP-pro-vWF to visualize secretion over time. **B,** Images of histamine and vehicle (DMSO)-treated cells between indicated groups. **C,** Quantification of vWF fluorescent intensity between indicated conditions. N= individual cells. White box denotes magnification; NS= not significant, values are means +/- SEM; n= individual sprouts; significance: **P<0.01, NS=Not Significant. Statistical significance was assessed with 1- way ANOVA followed by a Dunnett multiple comparisons test. All experiments were performed in triplicate.

**Supplemental Figure 6.**
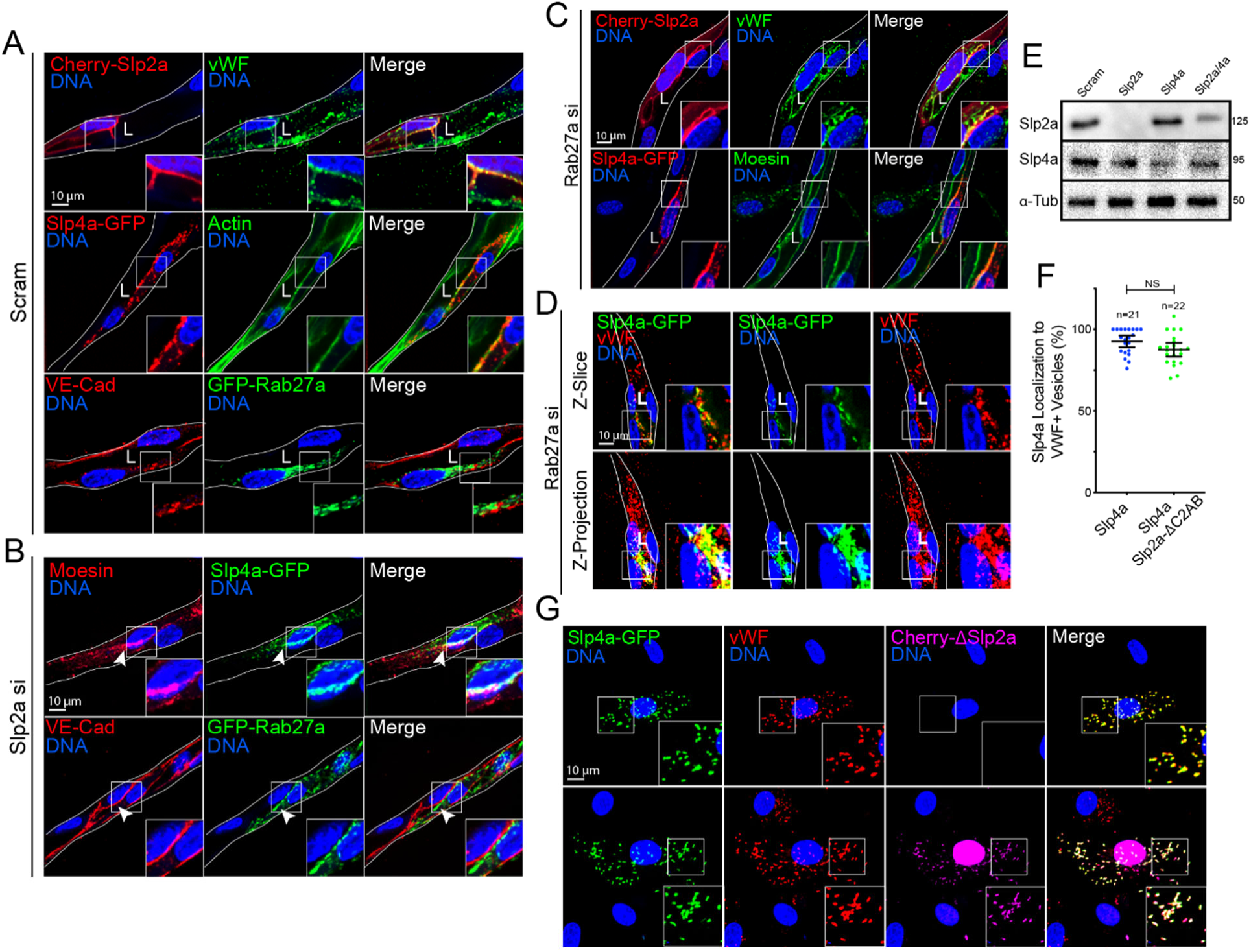
Slp2a and Rab27a do not affect each other’s localization during lumen formation. **A**, Scramble (scram) siRNA(si) knockdown sprouts showing localization of mCherry(Cherry)-Slp2a, GFP-Slp4a, and GFP-Rab27a. Sprouts were stained for von-Willebrand factor (vWF), actin and VE-cadherin (VE-cad). **B**, Slp2a siRNA knockdown sprouts expressing GFP-Slp4a and GFP-Rab27a probed for moesin and VE-cadherin. Arrowheads indicate a lack of lumen. **C**, Rab27a siRNA knockdown sprouts expressing mCherry-Slp2a and Slp4a-GFP stained for vWF and moesin. **D**, Rab27a siRNA knockdown sprouts expressing GFP-Slp4a and stained for vWF. The top row shows a single z-slice of the sprout and the bottom row shows a z-projection. **E,** Western blot confirmation of effects of Slp2a siRNA treatment on Slp4a expression as well as the reciprocal, Slp4a siRNA effects on Slp2a expression. n=3. **F,** Quantification of Slp4a localization intensity to vWF-positive vesicles with and without Cherry-Slp2a-ΔC2AB coexpression. **G,** Localization of Slp4a to vWF-positive vesicles with and without expression of cherry-Slp2a-ΔC2AB. In all panels L denotes lumen; white box denotes magnification; white lines denotes exterior of sprout, NS denotes Not-Significant. NS= not significant, values are means +/- SEM; n= individual sprouts; significance: Statistical significance was assessed with unpaired t- test.

**Supplemental Figure 7.**
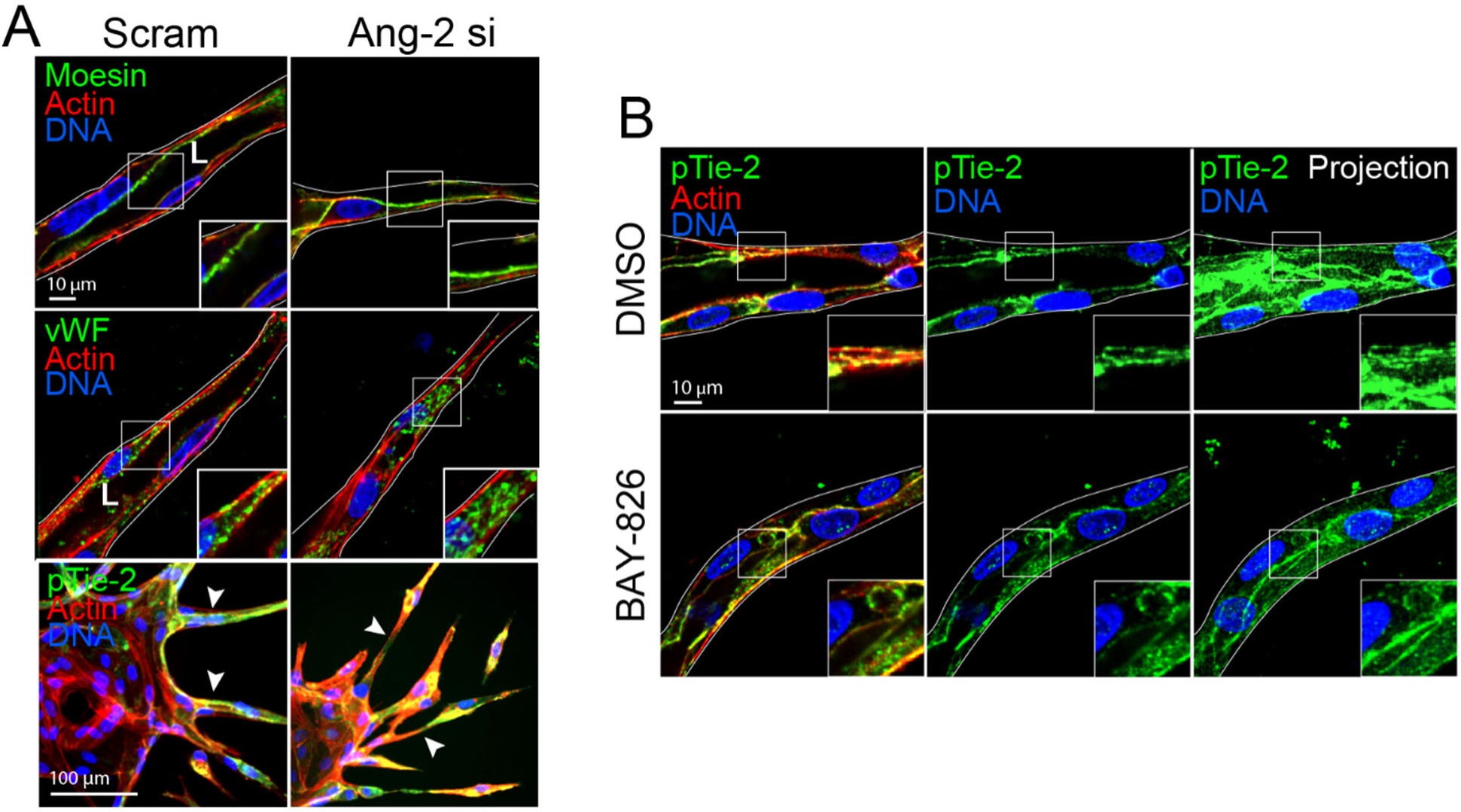
Ang-2 and Tie-2 inhibition promotes lumen defects. **A**, Scramble (scram) control and angiopoietin-2 (Ang-2) siRNA (si) knockdown sprouts stained for moesin, von Willebrand factor (vWF) and phosphorylated Tie-2 (pTie2). **B**, DMSO and BAY-826 (Tie-2 small molecule inhibitor) treated sprouts. Projections show level of pTie-2 staining compared with DMSO control. All experiments use human umbilical vein endothelial cells. In all panels L denotes lumen; white box denotes magnification; white lines denotes exterior of sprout. All experiments were performed in triplicate.

**Supplemental Figure 8.**
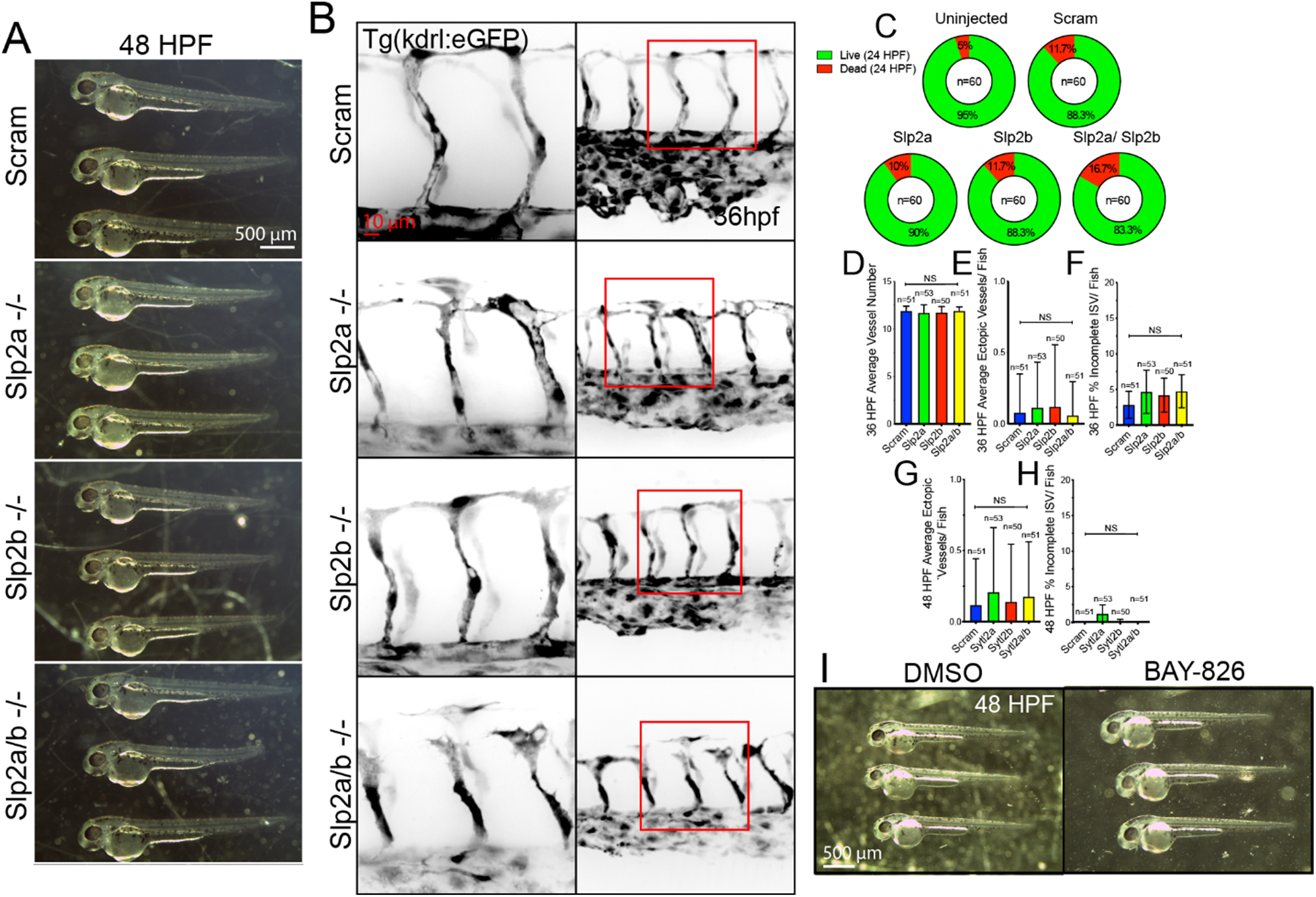
Slp2a/b KO zebrafish defects at 36 hpf and whole embryo morphology. **A,** Whole embryo morphology at 48 hours post fertilization (hpf) for indicated conditions. **B**, 36 hpf zebrafish intersomitic vessels (ISVs) for indicated conditions. Red boxes are area of higher magnification. **C,** Injection death rate at 24 hpf. **D,** Quantification of intersomitic vessel (ISVs) number at 36 hpf. **E,** Quantification of number ectopic vessels per fish at 36 hpf. **F,** Quantification of percent incomplete ISVs per fish at 36 hpf. **G**, Quantification of ectopic vessels per fish at 48 hpf. **H,** Quantification of percent incomplete ISVs per fish at 48 hpf. **I**, Representative image of 48 hpf larvae treated with DMSO (vehicle) or Tie-2 inhibitor Bay-826. NS= not significant, values are means +/- SEM; n= individual sprouts; significance: NS=Not Significant. Statistical significance was assessed with 1-way ANOVA followed by a Dunnett multiple comparisons test. All experiments were performed in triplicate.

**Supplemental Figure 9.**
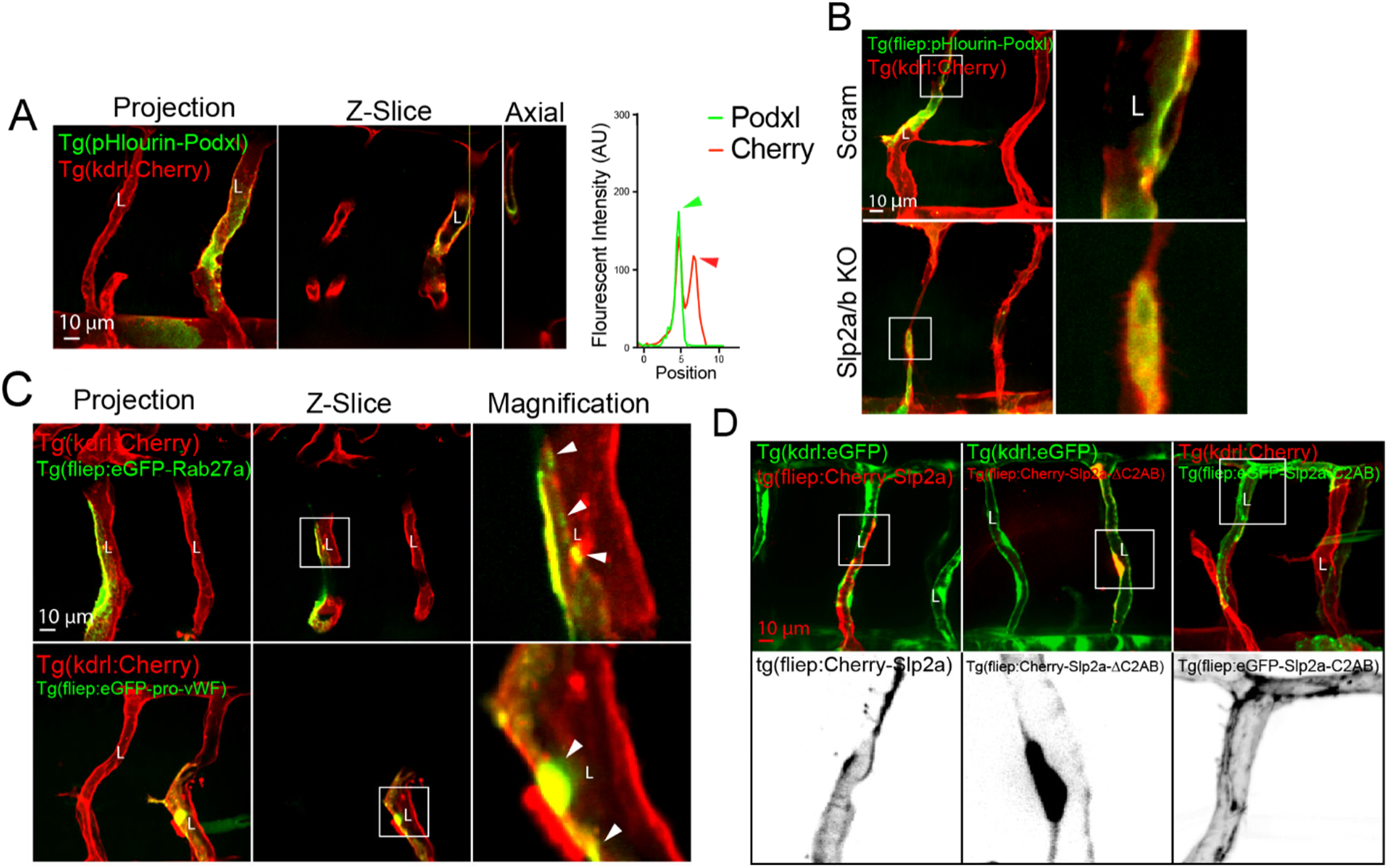
Visualization of lumen defects caused by Slp2a/b KO and localization of vWF, Rab27a and Slp2a. **A,** PHluorin-podocalyxin (Podxl) expression marking apical membrane in zebrafish intersomitc vessels (ISVs) on indicated background. Line scan depicts peaks in fluorescent intensity at the apical membrane. Green is apically localized pHluorin and red is endothelial mCherry (Cherry). **B**, PHlourin-Podxl localization between indicated scrambled (scram) and Slp2a/b CRISPR knockouts (KO). **C**, Mosaic expression of Human eGFP- Rab27a and pro-von Willebrand factor (vWF) in zebrafish ISVs. Arrowhead denote puncta. **D**, Zebrafish ISVs expressing Human mCherry(Cherry)-Slp2a, Cherry-Slp2a-ΔC2AB and Cherry-Slp2a-C2AB on a tg(kdrl:GFP) background. All zebrafish are at 48 hours post fertilization. In all panels L denotes lumen; white box denotes magnification.

